# Comodulation reduces interindividual variability of circuit output

**DOI:** 10.1101/2023.06.03.543573

**Authors:** Anna C. Schneider, Omar Itani, Elizabeth Cronin, Nelly Daur, Dirk Bucher, Farzan Nadim

## Abstract

2

Ionic current levels of identified neurons vary substantially across individual animals. Yet, under similar conditions, neural circuit output can be remarkably similar, as evidenced in many motor systems. All neural circuits are influenced by multiple neuromodulators which provide flexibility to their output. These neuromodulators often overlap in their actions by modulating the same channel type or synapse, yet have neuron-specific actions resulting from distinct receptor expression. Because of this different receptor expression pattern, in the presence of multiple convergent neuromodulators, a common downstream target would be activated more uniformly in circuit neurons across individuals. We therefore propose that a baseline tonic (non-saturating) level of comodulation by convergent neuromodulators can reduce interindividual variability of circuit output. We tested this hypothesis in the pyloric circuit of the crab, *Cancer borealis*. Multiple excitatory neuropeptides converge to activate the same voltage-gated current in this circuit, but different subsets of pyloric neurons have receptors for each peptide. We quantified the interindividual variability of the unmodulated pyloric circuit output by measuring the activity phases, cycle frequency and intraburst spike number and frequency. We then examined the variability in the presence of different combinations and concentrations of three neuropeptides. We found that at mid-level concentration (30 nM) but not at near-threshold (1 nM) or saturating (1 μM) concentrations, comodulation by multiple neuropeptides reduced the circuit output variability. Notably, the interindividual variability of response properties of an isolated neuron was not reduced by comodulation, suggesting that the reduction of output variability may emerge as a network effect.

**Significance Statement:** Neuromodulation has been explored as a mechanism to provide flexibility to the output of neural circuits. All neural circuits are subject to neuromodulation by multiple substances. Here, we propose a different but complimentary role for neuromodulation. We use two general facts about neuromodulation to show that convergent comodulation at tonic mid-concentration levels reduces interindividual variability of neural circuit output. These facts are that 1) Multiple neuromodulators can have convergent actions on the same subcellular substrates and 2) Neurons express different neuromodulator receptors at different levels.

## 4 Introduction

Historically, interindividual differences in animal behavior and in the function of neural circuits that underlie these behaviors have been disregarded, and most studies have examined the average behavioral response and the mean activity of the neural circuits involved (Asahina et al. 2022). Yet, recent studies have shown that behaviors as simple as reflexes or as complex as physiological responses to psychedelics can vary significantly among individuals (Cerins et al. 2022; Moujaes et al. 2022; Xu et al. 2022), even when accounting for genotype and other variables (Hageter et al. 2021; Palavicino-Maggio and Sengupta 2022; Rihani and Sachse 2022). Behavioral variability results largely from variability of underlying neural circuits (Feierstein et al. 2015; Rihani and Sachse 2022; Tamvacakis et al. 2022) and such variability is present even in smaller invertebrate neural circuits (Goaillard et al. 2009; Goaillard and Marder 2021; Hamood and Marder 2014). Despite significant variability, a neural circuit can generate an output that is good enough to produce a related behavior (Nassim 2018). Yet, in most cases, the CNS precisely controls behavior, and significant variability results in outputs deemed dysfunctional. Thus, at some level, circuit output must become constrained enough to produce consistent and meaningful behavior.

Several factors have been proposed to promote a consistent output across individuals at the level of an individual neuron. These include correlations of ion channel expression (Golowasch et al. 2017; Khorkova and Golowasch 2007; Temporal et al. 2011; Tobin et al. 2009; Tran et al. 2019), output degeneracy (Goaillard and Marder 2021; Ransdell et al. 2013), and excitatory neuromodulation (Schneider et al. 2022). Whether any of these mechanisms result in reduction of interindividual variability at the circuit level remains mostly unexplored. Here, we propose that convergent comodulation could play an important role in ensuring that neural circuits produce consistent circuit output in the face of extensive interindividual variability (Grashow et al. 2009; Maloney 2021).

Neuromodulation is conventionally thought to provide flexibility in neural circuit operation (Nadim and Bucher 2014). Yet, at any time, all neural circuits are comodulated by multiple substances (Harris-Warrick 2011; Marder 2012; Russo 2017). Some modulators control various aspects of excitability and synaptic output, and may target different neuron types (Harris-Warrick 2011). Others have limited but consistent actions across circuit neurons (Nadim and Bucher 2014). Consequently, different modulators often share subcellular targets, while remaining distinct in receptor expression levels or circuit neuron targets. Based on these facts, we propose an additional role for neuromodulation: that baseline (tonic) comodulation by multiple neuromodulators that have convergent effects, but different neural targets or patterns of receptor expression, is essential for producing consistent circuit output.

We tested this hypothesis in the pyloric network of the stomatogastric ganglion (STG) of the crab *Cancer borealis*, which has been a long-standing testbed for neuromodulation (Daur et al. 2016; Marder et al. 2014; Stein 2009). The pyloric circuit consists of identified neurons with known synaptic connections. Under physiological conditions it expresses regular periodic activity (∼1 Hz) that can be easily quantified. Upon removal of all neuromodulatory inputs (decentralization) the pyloric rhythm is disrupted or significantly slowed and shows significant interindividual variability (Hamood et al. 2015). We measured several circuit activity attributes (cycle frequency, activity phases, etc.) after decentralization, and in the presence of one, two, or three neuropeptides, applied at the same total concentration. We then quantified the effect of comodulation on the interindividual variability of these attributes. We also used the same peptide combinations to assess the interindividual variability of the activity attributes of a single isolated pyloric neuron that is targeted by all three neuropeptides to see if comodulation of circuit variability is consistent with the effect of variability at the single-cell level. As a final step, we examined if a minimally constrained family of pyloric circuit model neurons captures the experimental observations on the effects of comodulation on interindividual variability.

## 5 Methods

### 5.1 Experimental preparation

Experiments were performed on the isolated stomatogastric nervous system (STNS) of male Jonah crabs (*Cancer borealis*) obtained from local fish markets and maintained in recirculating artificial seawater tanks at 12°C with a 12:12 hour light-dark cycle until use. Crabs were anesthetized by chilling in ice for at least 30 min before removing the stomach. The STNS was carefully dissected from the stomach and pinned out in a Sylgard (Dow Corning) lined petri dish. All STNS preparations consisted of the stomatogastric ganglion (STG), the anterior portion that contains modulatory projection neurons, and motor nerves including the lateral ventricular (lvn), pyloric dilator (pdn) and pyloric constrictor (pyn) nerves (Fig. 1A). The sheath around the STG was removed with fine tungsten pins to facilitate intracellular recording and the penetration of bath applied chemicals. A large petroleum jelly well was constructed around the STG to allow for fast exchange of solutions and was constantly perfused with cold saline (11-12°C). Smaller wells for extracellular recordings were constructed around the lvn, pdn, and pyn. To remove the influence of neuromodulatory projection neurons on the STG neurons, the stomatogastric nerve (stn) was transected with fine scissors (decentralization).

**Figure 1:**
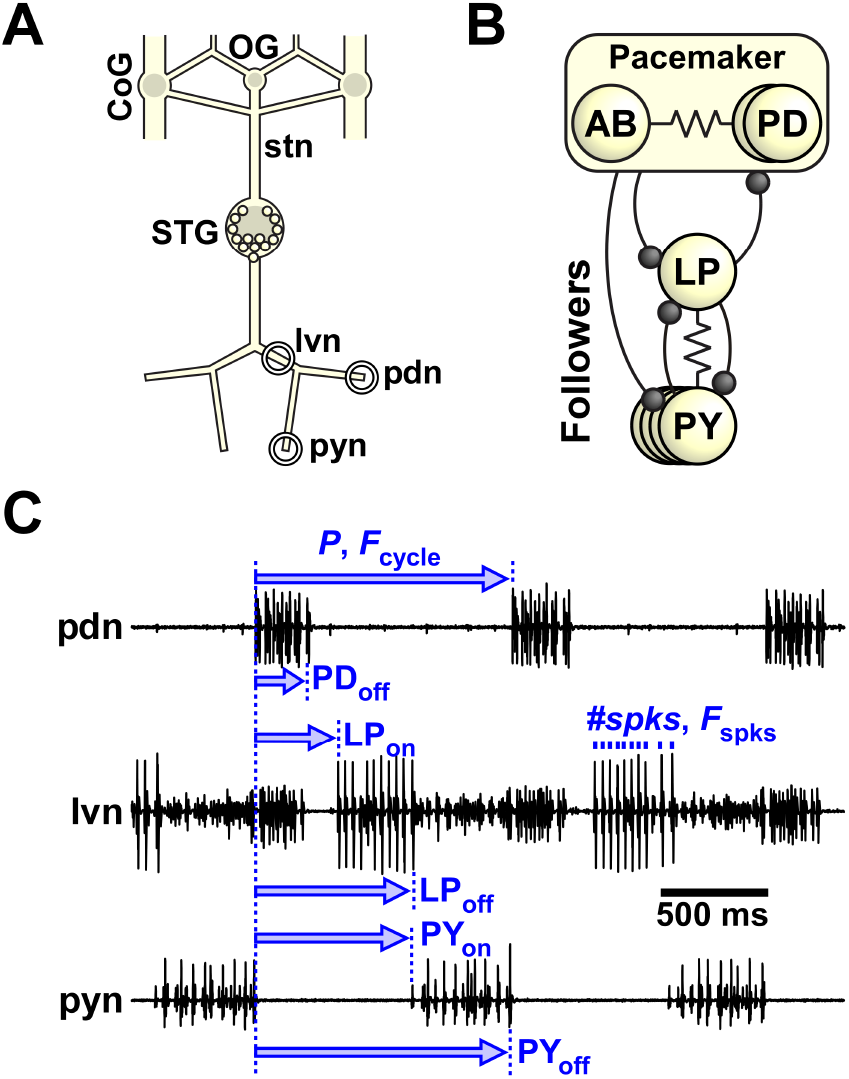
The stomatogastric nervous system (STNS) of crabs. **(A)** Schematic of the STNS of the crab *C. borealis*. Neurons of the pyloric circuit are located in the stomatogastric ganglion (STG). They receive input from neuromodulatory projection neurons that originate in the paired commissural ganglia (CoG) and the unpaired esophageal ganglion (OG) and send their axons through the stomatogastric nerve (stn). Pyloric motor neurons project their axons through the lateral ventricular (lvn), pyloric dilator (pdn) and pyloric constrictor (pyn) nerves. The extracellular recording sites are indicated by circles. **(B)** The core pyloric circuit consists of a group of pacemaker neurons: The anterior burster (AB) and two pyloric dilator (PD) neurons. They inhibit their followers, the lateral pyloric (LP) and 3-5 pyloric constrictor (PY) neurons. Inhibitory synapses are shown as circles, electrical connections are depicted with resistor symbols. **(C)** The pyloric rhythm and rhythm parameters. The activity of the PD neurons from the pacemaker group is recorded from the pdn. Activity of all neurons can be recorded from the lvn, where LP is typically the largest unit, PD is mid-sized, and PY are the smallest units. In addition, PD and PY activity are also recorded from the pdn and pyn, respectively. Cycle period is defined as the time from the beginning of one PD burst (PD on) to the next. *F*_cycle_ is the inverse of cycle period (*P*). Burst start (on) and end (off) for each neuron is calculated by dividing the latency with respect to PD on by the cycle period. In addition, we counted the number of spikes per burst (#spks) and calculated the average spike frequency (F_spks_) by dividing #spks-1 by burst duration (off-on).

### 5.2 Solutions

*C. borealis* saline contained (in mM): 440 NaCl, 26 MgCl_2_, 13 CaCl_2_, 11 KCl, 10 Tris base, 5 maleic acid, buffered to pH 7.4. Proctolin (PROC; RS Synthesis), crustacean cardioactive peptide (CCAP; RS Synthesis), and red pigment concentrating hormone (RPCH; GenScript) were prepared as 10^−3^ M aliquots (PROC and CCAP in distilled water, RPCH in dimethyl sulfoxide: DMSO, Fisher Scientific) and stored at −20°C until use. The stock solutions were thawed and diluted to a final concentration in either normal saline, or saline with picrotoxin (PTX, Sigma-Aldrich), immediately before the experiments. The final total modulator concentrations were 10^−9^ M (low), 3×10^−8^ M (mid), and 10^−6^ M (high). In a few initial experiments, a concentration of 10^−8^ M was used as mid. The total modulator concentration (low, mid, or high) was held constant in comodulator applications so that with two modulators each was 1/2 of the total concentration and with three modulators each was 1/3 of the total concentration.

For experiments that assessed single neuron excitability, synapses were blocked with PTX. PTX was dissolved in DMSO at 10^−2^ M and stored as stock solution at 4°C. Immediately before the experiment, PTX stock solution was diluted to a final concentration of 10^−5^ M in saline.

### 5.3 Electrophysiology

Thin stainless steel pin electrodes were placed inside and outside petroleum jelly wells around the lvn, pdn, and pyn to record the pyloric rhythm extracellularly. The extracellular electrodes were connected to a differential AC amplifier (Model 1700, A-M Systems). To reach a steady state in each neuromodulatory condition, we waited 20-30 minutes after decentralization before washing in any neuromodulator, and 10-15 minutes before switching from one neuromodulator cocktail to the next. For analysis, we recorded the pyloric rhythm for 1-2 minutes each in the intact STNS, 20 min after decentralization, and 10 min after washing in each of the sequence of neuropeptides applied at different concentrations and combinations. In total, we obtained four different datasets, each with intact and decentralized conditions in addition to: 1) low PROC (P), low PROC + CCAP (PC), low PROC + CCAP + RPCH (PCR), mid P, mid PC, mid PCR, wash out; 2) mid P, mid PC, mid PCR, wash out 3) high P; 4) high PC. Because modulators were added sequentially, we did not wash out with saline between modulatory conditions. To minimize the variability that can be caused by changing environmental or experimental factors we typically did these experiments simultaneously with two preparations in the same dish and at the same time of the day.

To examine the variability of intrinsic response properties, we used intracellular two-electrode current clamp recordings. Intracellular voltage recording and current injection were done with Axoclamp 900A amplifiers (Molecular Devices). All recordings were digitized at 5 kHz (Digidata 1440A, Molecular Devices) and recorded with Clampex 10.6 (Molecular Devices).

We first identified the lateral pyloric (LP) and the two pyloric dilator (PD) neurons in an intact preparation by matching the intracellular activity to the extracellular activity on the lvn and pdn, and by their characteristic intracellular membrane potential waveforms. We then decentralized the preparation, impaled the LP neuron with two electrodes and both PD neurons with one electrode each, and washed in PTX for at least 15 minutes until inhibitory postsynaptic potentials in the PD neuron (from the presynaptic LP neuron) disappeared and LP inhibition during PD bursts was greatly reduced. We then ran current clamp protocols as described in Schneider et al. (2022) in the LP neuron while hyperpolarizing the PD neurons with −5 nA DC current injection to eliminate the remaining PTX-insensitive PD to LP synaptic inhibition. Each set of protocols was repeated in decentralized, mid P, mid PC, mid PCR, and wash. To obtain *f-I* curves, we depolarized LP by injecting current steps from 0 to 5 nA, in increments of 0.5 nA. To measure *f-I* hysteresis, we also used the inverse sequence, from 5 nA to 0 nA. We ran each protocol twice and used the average spike responses for analysis (see below). To measure rebound properties, we used two protocols. First, we hyperpolarized LP five times with 10 s current injections of −5 nA, interspersed with 10 s recovery time. This interval was long enough for LP to return to initial conditions before the next sweep (Schneider et al. 2022). Second, we hyperpolarized LP periodically with twenty 1 s-on/1 s-off, −5 nA current pulses to mimic more realistically the LP inhibition by the PD neurons. To analyze the response to periodic hyperpolarization, we only used the last 10 pulses to avoid the transient responses of the LP neuron to the first few pulses (Schneider et al. 2022).

### 5.4 Data analysis

All data were imported from Clampex to Matlab (version 2021a) using the “abfload” function (Hentschke 2011), and analyzed with custom written scripts and the “CircStat” toolbox (Berens 2009). Statistical tests were performed with SigmaPlot (version 12.0, Systat Software) with a significance level of α = 0.05. We used ANOVA with Tukey’s post-hoc test if the data passed tests for normal distribution (Shapiro-Wilk) and equal variance (Levene). If one of these tests failed, we used ANOVA on ranks with Dunn’s post-hoc test to compare metrics between different modulatory conditions.

#### Pyloric rhythm attributes

To quantify the pyloric rhythm (Fig. 1), we used extracellular nerve recordings to calculate the cycle period (*P* = time difference between the onsets of two consecutive PD bursts), and, for PD, LP, and pyloric constrictor (PY) each, the latencies of burst onset (e.g., LP_on_) and termination (e.g., LP_off_) relative to PD burst onset, number of spikes per burst (#spks), and mean intra-burst spike frequency (*F*_spks_). Cycle frequency (*F*_cycle_) was calculated as 1/*P*, and the burst onset and end phases were calculated as the respective latency divided by cycle period. For example, the end phase of the LP burst is given by ϕLP_off_ = LP_off_ / *P*. Note that in each animal, PD exists in two copies and PY in 3-5 copies. Since we obtained #spks and *F*_spks_ from extracellular recordings, the values we report were from all active PD or PY units, detected on pdn and pyn, respectively. It is worth noting that we used bursting activity phases, rather than latencies, as important attributes to measure. The reason for this is that burst latencies, but not phases, show a strong correlation with *F*_cycle_ as has been noted in a number of publications (e.g., Anwar et al. 2022; Bucher et al. 2005; Cronin et al. 2023; Goaillard et al. 2009). Similarly, we used both *#spks* and *F*_spks_ because they only weakly correlated in the pyloric circuit (Bucher et al. 2005; Cronin et al. 2023).

A complete quantification of the pyloric circuit activity requires the existence of a triphasic rhythm. In our data summaries we indicate the number of preparations of each dataset that were analyzed. The missing preparations did not express rhythmic activity, which sometimes occurs in decentralized unmodulated preparations.

We used different metrics to measure the interindividual variability of the different attributes. For *F*_cycle_, *#spks*, and *F*_spks_, we used the adjusted coefficient of variation (CV), which is the standard deviation divided by the mean and adjusted for sample size (Haldane 1955). Since CV only is meaningful for data on a ratio scale, we used the circular variance as a measure of variability for φ. Briefly, each value of φ is represented as a vector composed of the sine and cosine of its angle on a unit circle with a length of 1. Averaging these vectors gives a resultant vector with the mean φ as direction and a length (r) between 0 and 1 that depends on the spread of the data. If all values of φ data were similar, and therefore located at the same point on the unit circle, the r-vector would have a length of 1. If all data were evenly spread around the circle, r-vector length would be 0. Circular variance is, by definition, 1 – r. Since the circuit neurons do not have independent activity phases, we computed the correlation coefficient matrices for all phases for each modulatory condition and their eigenvalues. We then summed the eigenvalues, which yields a measure for the variability of all phase data.

#### Single neuron excitability attributes

We analyzed the current clamp data as described in Schneider et al. (2022). In brief, we fit the *f-I* curves with the power function

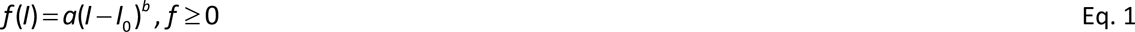

where *a* is a scaling factor approximating the maximum spike frequency, *I*_0_ is the (calculated) current level that first elicited spikes, and the power *b* (set to 0 ≤ *b* ≤ 1 to limit the *f-I* curve to a sublinear function) measures the nonlinearity of the *f-I* curve. Hysteresis was calculated as the ratio of the average spike frequency for injected current between 2 to 4 nA with increasing current injection divided by the average spike frequency between 2 to 4 nA with decreasing current injection.

For the rebound protocols, we considered all five sweeps for the 10 s current injection and only the last 10 sweeps for the 1 s current injection. We measured the average latency between the end of the current injection and the first spike. We fitted the cumulative spike histogram with the sigmoid

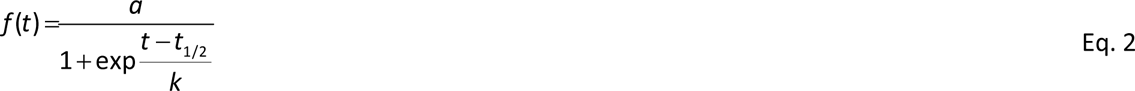

where *a* is the total number of spikes in the interval between current injections, *t*_1/2_ is the sigmoid midpoint time relative to the end of current injection, and *k* is the slope factor.

As a metric for interindividual variability of fit parameter, hysteresis, and latency, we used either the adjusted CV, or standard deviation for interval measures (for which CV was not applicable).

### 5.5 Modeling

#### Models of the pyloric circuit

Computational models based on the pyloric circuit were constructed with the circuit diagram given in Fig. 1, but with only one PD and 2 PY neurons. Models were constructed and run using the Python (ver. 3.7) (Van Rossum and Drake 2009) interface to NEURON (ver. 8.2.1) (Carnevale and Hines 2006). Pyloric neurons other than AB were modeled to have two compartments, one representing the soma, neurites and dendrites (S/N), and one representing the axon (A). The AB neuron was represented by a single somatic compartment, based on our previous model of the pacemaker group (Daur et al. 2019). Synaptic connections were modeled to follow sigmoidal transfer functions with activation and decay time constants depending on the presynaptic membrane potential. All model parameters are described in Table 1.

**Table 1:** Main components of the pyloric circuit model. Model equations are provided in the supplement Figure 10-1. The parameters for intrinsic currents and synapses are, respectively, in the supplements Figure 10-2 and Figure 10-3.

| Cells | Synapses | Intrinsic currents |
| --- | --- | --- |
| AB | AB → LP, PY1, PY2 (inhibitory) | leak |
| PD | PD → LP, PY1, PY2 (inhibitory) | Na, CaS, MI |
| LP | LP → PD, PY1, PY2 (inhibitory) | KS, A, Kd |
| PY1 | PY1, PY2 → LP (inhibitory) | |
| PY2 | AB ↔ PD (symmetric electrical coupling) | |
| | LP ↔ PY1, PY2 (rectifying electrical coupling) | |

#### Initial family of models

To build an initial family of pyloric circuit models, we started with a general form of the model as described above. We then allowed the maximal conductances of intrinsic currents and synapses to vary. We used the simulation-based inference python toolbox (SBI v0.20.0) to select 175 models that complied with attributes of activity within the following selection criteria:

- *F*_cycle_ > 0,
- φ: PD_off_ < 1; LP_on_ < 1; LP_off_ < 0.65; PY_on_ < 1; PY_off_ < 1.2,
- *#spks*: PD > 2; LP > 2; PY (1 & 2 combined) > 1.

To simulate decentralization (Hamood et al. 2015), we ran simulations of each model while scaling down the value of *g*_MI_ in all neuron types multiplicatively until the model pacemakers (AB and PD) ceased to produce bursting oscillations. We chose the values of *g*_MI_ at the cusp of pacemaker activity as our decentralized case to allow for quantifiable rhythms.

#### Modeling peptide comodulation

We modeled peptide modulation by increasing the maximal *g*_MI_ values multiplicatively across each cell type. To capture multiple axes of peptide modulation, we scaled *g*_MI_ along three axes: pacemakers (AB/PD), LP, and PYs, resulting in a 3D grid spanning multiplicative scalars from 1 to 1.75 in increments of 1/16. Across the 175 models, we measured interindividual variability of each attribute at each point in the grid using circular variance (for φ) or CV (all other attributes) and reported it as percent change with respect to the variability of the decentralized case.

## 6 Results

### 6.1 Comodulation at mid concentrations reduced interindividual variability of the circuit activity attributes

The intact stomatogastric nervous system consists of the stomatogastric ganglion (STG), which includes the pyloric circuit neurons, as well as the esophageal ganglion (OG) and the paired commissural ganglia (CoGs; Fig. 1A). A variety of small-molecule neurotransmitters and neuropeptides modulate circuit activity in the STG, either as neurohormones present in the hemolymph or released from descending projection neurons. These projection neurons, many of which are spontaneously active at any time, have their cell bodies in the OG and CoGs and send axons to the STG via the stomatogastric nerve (stn). We characterized the pyloric rhythm by determining output pattern attributes from simultaneous extracellular recordings of the three motor nerves, lvn, pdn and pyn (Fig. 1A). The core circuit that generates the pyloric rhythm (Fig. 1B) includes the anterior burster (AB) interneuron as well as two pyloric dilator (PD), lateral pyloric (LP) and 3-5 pyloric constrictor (PY) motor neurons. The axons of the three motor neuron types, PD, LP and PY, project bilaterally through the lvn and their action potentials can be readily separated by simultaneous extracellular recordings of lvn, pdn and pyn that show the well-characterized triphasic pyloric burst sequence of these neurons (Fig. 1C)(Marder and Bucher 2007).

We measured pyloric output pattern attributes (marked in Fig. 1C) across different modulatory states, with intact descending input, after decentralization (see below), and after application of neuropeptides (Fig. 2A). With intact descending inputs, spontaneous activity of projection neurons ensures that the pyloric neurons are always bathed in a soup of modulatory neurotransmitters, including several peptides that are co-released by these projection neurons (Nusbaum et al. 2017). In vitro, all modulatory inputs can be removed by blocking or severing the projection nerve (stn), a procedure we refer to as decentralization (Fig. 2A). Removing all neuromodulatory input to the STG by decentralization greatly slows or even disrupt the pyloric rhythm (Hamood and Marder 2015). Bath application of excitatory neuropeptides have been claimed to produce a peptide-specific version of the pyloric rhythm (Marder and Thirumalai 2002; Marder and Weimann 1992; Swensen and Marder 2001). We used three excitatory neuropeptides that target distinct but overlapping subsets of neurons in the pyloric circuit (Fig. 2A), but that all converge on the same target ion channel within these subsets. This target is a voltage-gated inward current referred to as *I*_MI_ (Golowasch and Marder 1992; Gray et al. 2017; Gray and Golowasch 2016; Schneider et al. 2021; Swensen and Marder 2000). PROC activates *I*_MI_ in AB, PD, LP, and most PY neurons, while both CCAP and RPCH activate *I*_MI_ only in AB and LP neurons (Fig. 2A) (Swensen and Marder 2001). Additionally, both PROC and CCAP enhance synaptic currents in the reciprocal connections between the pacemaker neurons and LP (Li et al. 2018), but effects on the pacemaker to PY and PY to LP synapses have not been studied. RPCH increases the strength of the LP to pacemaker synapse (Atamturktur and Nadim 2011; Thirumalai et al. 2006), but its effect on other synapses has not been studied.

**Figure 2:**
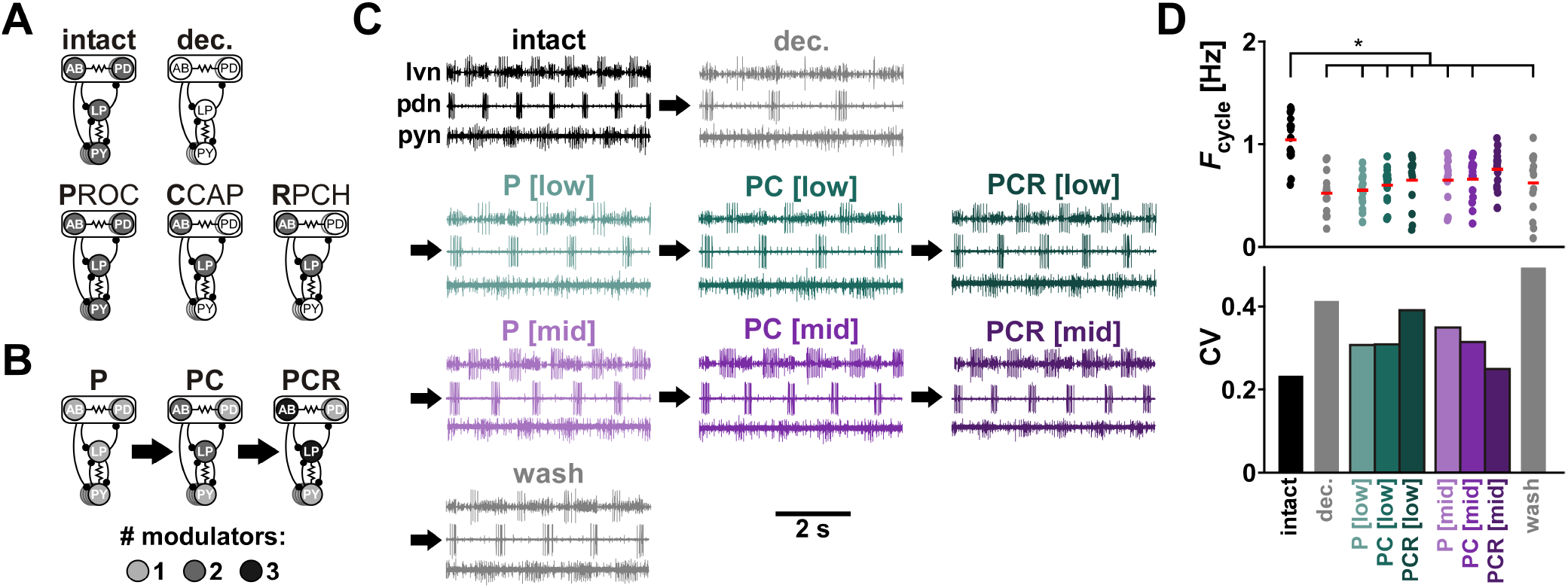
Comodulation at mid concentrations reduces the interindividual variability of cycle period. **(A)** Schematic indication of neuromodulator target neurons (filled circles) in the different neuromodulatory conditions. In the intact condition, each neuron is targeted by an unknown number of neuromodulators. After decentralization (transection of the stomatogastric nerve), all neuromodulation is removed. Proctolin (PROC, P) targets all pyloric neurons, whereas CCAP (C) and RPCH I target only AB and LP neurons. **(B)** Overlapping targets of comodulation by PROC, CCAP, and RPCH. Shading indicates how many of the applied neuromodulators target each neuron of the pyloric circuit. **(C)** Extracellular recordings of the pyloric rhythm from one animal under different neuromodulatory conditions (color coded). After decentralization, we applied increasing numbers (P, PC, PCR) and increasing concentrations ([low]: 10^−9^ M, [mid]: 3 x 10^−8^ M) of neuropeptides (arrows), followed by washing out all neuromodulators. **(D)** *F*_cycle_ and the corresponding CV (standard deviation / mean) under different modulatory conditions. Individual dots represent data from individual experiments, red bars indicate the mean value. N = 15 animals. Asterisks indicate significant differences between groups (Dunn’s post-hoc test, p ≤ 0.05). Total modulator concentration for [low]: 10^−9^ M; [mid]: 3 x 10^−8^ M. ANOVA results in Table 2. Figure 2-1: Raw data for F_cycle_.

**Table 2:** ANOVA results for pyloric rhythm parameters at low and mid modulator concentrations. If tests for normality or equal variance failed, results are for ANOVA on ranks. Groups are intact, decentralized, P [low], PC [low], PCR [low], P [mid], PC [mid], PCR [mid], wash. p values smaller than α are printed in bold. N: number of animals; df: degrees of freedom; res: residual. Note that, despite the significant effect from ANOVA, the pairwise comparison for PY on phase did not show any significant differences between groups (Fig. 3A).

| N | parameter | normality | variance | test statistic | df | res. | p |
| --- | --- | --- | --- | --- | --- | --- | --- |
| 15 | Cycle frq | fail |  | H = 36.645 | 8 |  | **<0.001** |
| 15 | PD off | fail |  | H = 24.794 | 8 |  | **0.005** |
| 15 | LP on | fail |  | H = 19.799 | 8 |  | **0.011** |
| 15 | LP off | fail |  | H = 37.006 | 8 |  | **<0.001** |
| 15 | PY on | fail |  | H = 20.498 | 8 |  | **0.009** |
| 15 | PY off | fail |  | H = 15.495 | 8 |  | 0.050 |
| 15 | PD spikes | pass | pass | F = 3.365 | 8 | 119 | **0.002** |
| 15 | PD spike frq | fail |  | H = 29.773 | 8 |  | **<0.001** |
| 14 | LP spikes | pass | pass | F = 21.217 | 8 | 111 | **<0.001** |
| 14 | LP spike frq | pass | fail | H = 50.238 | 8 |  | **<0.001** |
| 15 | PY spikes | fail |  | H = 9.413 | 8 |  | 0.309 |
| 15 | PY spike frq | fail |  | H = 9.958 | 8 |  | 0.268 |

Our goal was to test whether comodulation by multiple excitatory neuropeptides reduces interindividual variability of circuit output. To examine this hypothesis, we bath applied combinations of one (PROC: P), two (PROC + CCAP: PC), and three (PROC + CCAP + RPCH: PCR) neuropeptides at the same total low and mid concentrations (Fig. 2B, C) in several preparations and examined the variability of multiple circuit output attributes (see Materials and Methods for detailed description). Statistical results for all pyloric circuit attributes of this dataset are listed in Table 2. Raw data are provided in Figure 2-1 and Figure 3-1.

**Figure 3:**
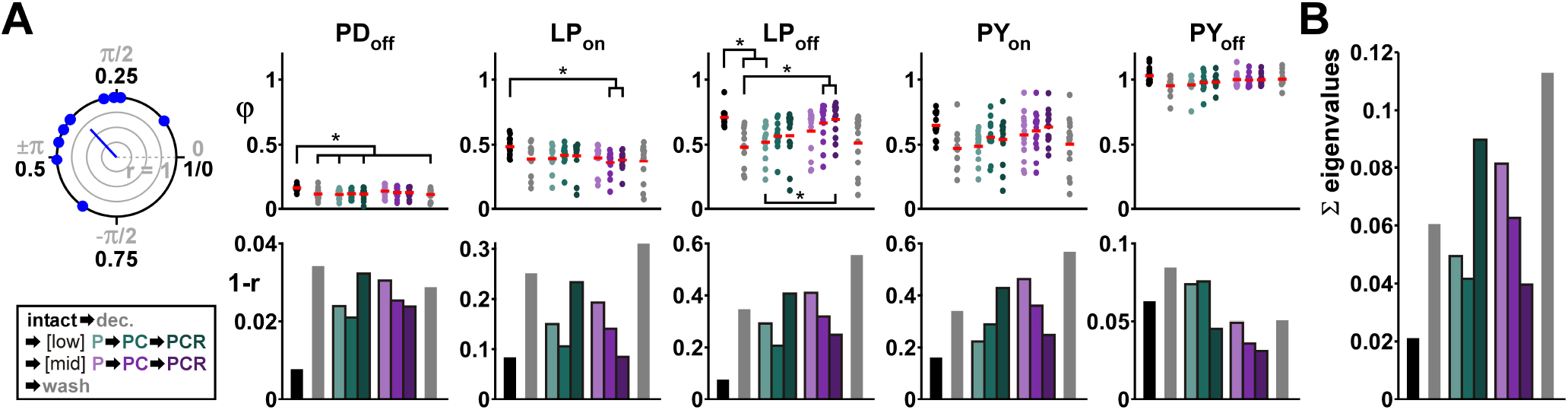
Comodulation at mid concentrations reduces the interindividual variability of pyloric rhythm parameters on the circuit output level. **(A)** Burst start (on) and termination (off) and the corresponding circular variance (1-r, see Methods) under different modulatory conditions (color coded). Individual dots represent data from individual experiments, red bars indicate the circular mean. INSET shows a circular plot of a randomly generated phase dataset (N = 10, x̅ = 0.35, µ = 0.15). The length of the r-vector indicates the spread of the data. Markers inside the circle indicate 25%, 50%, and 75% of the radius. **(B)** Sum of eigenvalues from the covariance matrix of the phases for each modulatory condition. N = 15 animals. Asterisks indicate significant differences between groups (Dunn’s post-hoc test, p ≤ 0.05). Total modulator concentration for [low]: 10^−9^ M; [mid]: 3 x 10^−8^ M. ANOVA results in Table 2. Figure 3-1: Raw data for panel A.

Decentralization decreased *F*_cycle_ compared to intact preparations. While application of one or more neuropeptides at low or mid concentrations to the decentralized preparation tended to increase *F*_cycle_, it was only with application of three modulators at mid concentration that cycle it recovered to values statistically similar to those in the intact condition (Fig. 2D, upper panel). Upon decentralization, the *F*_cycle_ coefficient of variation (CV) approximately doubled compared to the intact state and remained high when we applied one, two, or three neuropeptides at low concentration (Fig. 2D, lower panel). However, when we applied increasing numbers of peptides at mid concentration, the CV consistently decreased toward its intact value. Washing out the modulators increased CV as expected. A similar effect was observed for the on and off phases of all three neurons, PD, LP, and PY (Fig. 3). The burst onset (on) and end (off) phases of individual pyloric neurons within each cycle are a major determinant of the proper function of this central pattern generator, and these activity phases remain surprisingly consistent across individual animals despite large variations in *F*_cycle_ (Anwar et al. 2022; Bucher et al. 2005; Cronin et al. 2023; Goaillard et al. 2009). Decentralization and application of neuropeptides affected the mean phases of PD and LP, but not of PY (Fig. 3A top panels). However, there was a large effect on the interindividual variability of the phase values. We measured variability of activity phases as the circular variance (1 - r; Fig. 3A top left inset; also see Methods). Circular variance of all phases greatly increased upon decentralization and was somewhat reduced by application of neuropeptides (Fig. 3A bottom panels). Notably, for all phases, circular variance consistently decreased when we increased the number of applied neuropeptides at mid concentration but not at low concentration (Fig. 3A bottom panels).

Because of recurrent connectivity of the pyloric circuit (Fig. 1A), the PD, LP and PY on and off phases are not independent of each other. To account for the covariation of the phases, we computed the covariance matrix of all five phases in each modulatory condition. Total variation can be measured as the trace (equivalently, sum of the eigenvalues) of this matrix (Fig. 3B). The resulting pattern of total variation of phase under the different modulatory conditions was the same as for *F*_cycle_ and for individual phases: Variability increased with decentralization but consistently decreased when we increased the number of neuropeptides at mid concentrations. No consistent pattern was seen at low concentrations.

We also quantified two other important output attributes of the pyloric circuit neurons: #*spks* and *F*_spks_. In PD, #*spks* was largely unaffected by decentralization and modulator application (Fig. 4A top left). In contrast, decentralization decreased the PD *F*_spks_, and only mid concentrations peptide application returned this attribute toward its intact values (Fig. 4B top left). LP generated significantly more #*spks* in PC and PCR at mid concentrations (Fig. 4A top middle). This was also reflected in the LP *F*_spks_ (Fig. 4B top middle). In contrast, mean spiking activity of PY remained unchanged across all conditions.

**Figure 4:**
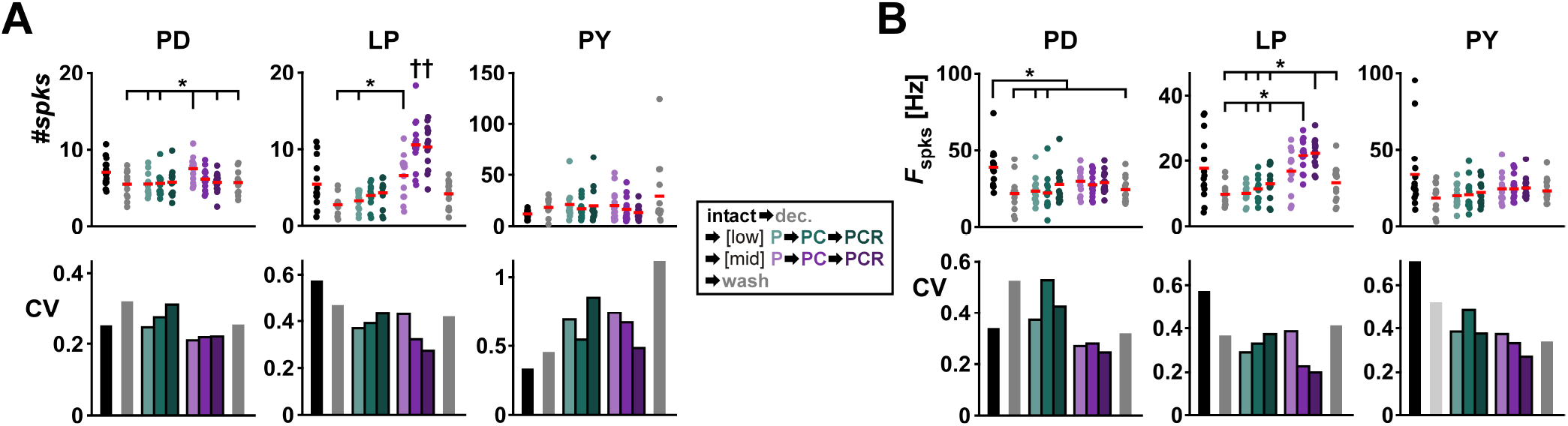
Comodulation at mid concentrations reduces the interindividual variability of burst parameters on the circuit output level. **(A)** Average number of spikes (#spks) per burst and corresponding CV for each type of neuron at each modulatory condition (color coded). **(B)** Average spike frequency (F_spks_) within a burst and corresponding CV for each type of neuron at each modulatory condition (color coded). N = 15 animals. Dots represent values from individual experiments, horizontal bars indicate the mean value. Asterisks indicate significant differences between groups (Dunn’s post-hoc test, p ≤ 0.05). Dagger indicates that this group is different from all others, except other groups marked with dagger. Total modulator concentration for [low]: 10^−9^ M; [mid]: 3 x 10^−8^ M. ANOVA results in Table 2. Figure 4-1: Raw data for panels A and B.

The changes in interindividual variability of the spiking activity (measured as CVs) were somewhat different from the consistent changes we saw for *F*_cycle_ and the activity phases. For instance, the CVs of PD #*spks* and *F*_spks_ did not consistently decrease with adding modulators at mid concentration but rather were similar across those conditions. However, in LP and PY, variability still decreased with additional modulators at mid concentration (purple bars in Fig. 4A, B, bottom left). Additionally, for both the LP and PY neurons, the CVs of both spiking attributes were reduced by adding modulators at mid concentration (purple bars in Fig. 4A, B, bottom middle & right).

One caveat of these experiments was that, because we had applied the neuropeptide sequentially at low and then mid concentrations (see, e.g., Fig. 2C), the preparations recorded with mid concentration of P, PC and PCR had all been pre-exposed to low concentrations of all three modulators (low PCR). To ensure that the effects of comodulation seen at mid concentration were not due to history-dependence, we repeated these experiments by applying P, PC and PCR at only mid concentrations. These experiments effectively replicated the result that most attributes of the pyloric circuit output showed a reduction of variability with increased numbers of peptide comodulators (Fig. 5, Table 3; raw data are provided in Figure 4-1). Once again, the CVs of PD spike attributes did not decrease with comodulation. In these experiments, we also did not see a consistent reduction in variability of PY spike attributes or end phase in the transition from two (PC) to three (PCR) modulators.

**Figure 5:**
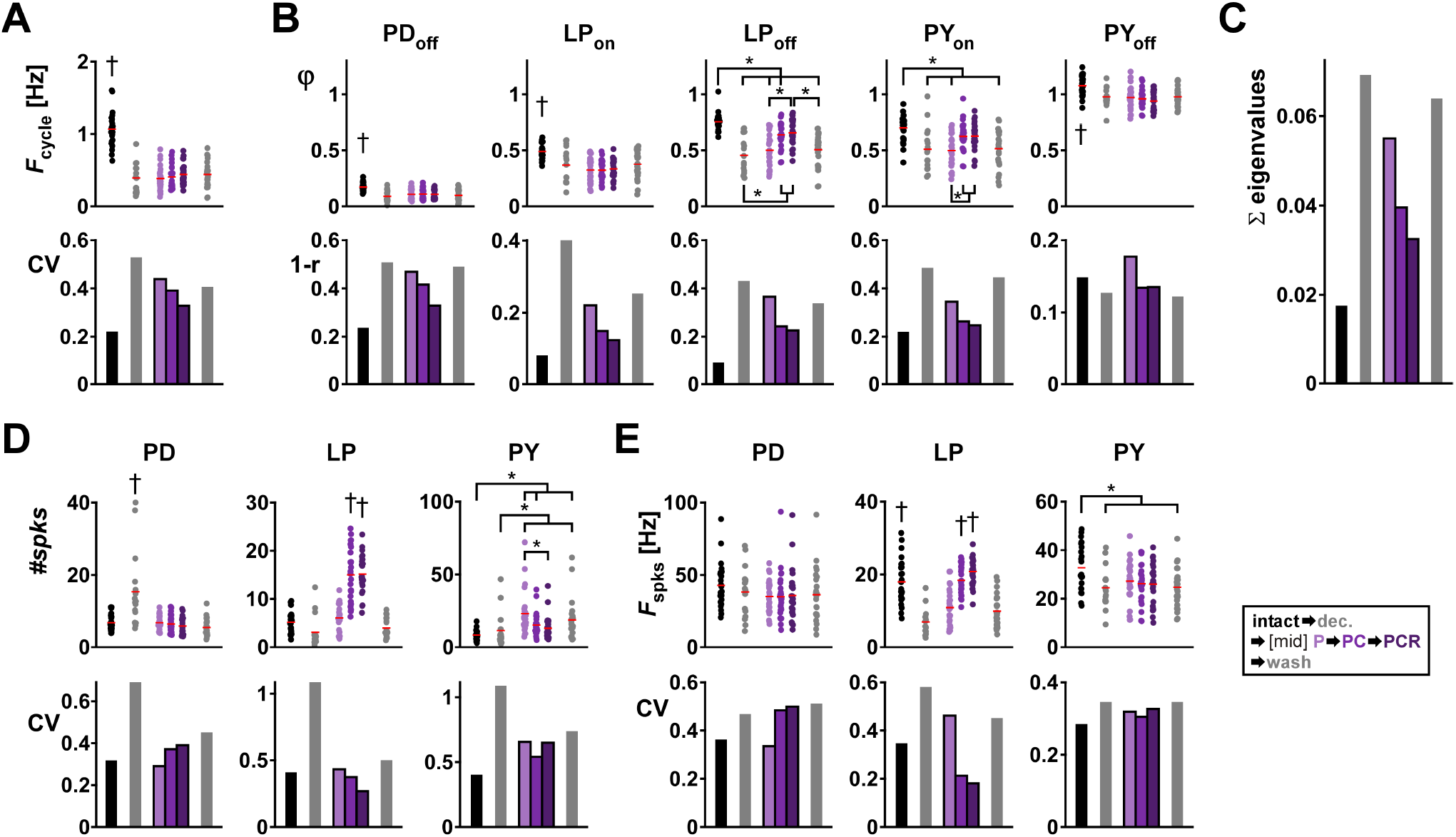
Comodulation at mid concentrations reduces the interindividual variability of rhythm parameters on the circuit output level in an independent dataset. **(A)** *F*_cycle_ and the corresponding CV under different modulatory conditions. **(B)** Burst start (on) and termination (off) and the corresponding circular variance, 1 - *r*, see Methods) under different modulatory conditions (color coded). **(C)** Sum of eigenvalues from the covariance matrix of the phases for each modulatory condition. **(D)** Average number of spikes (#spks) per burst and corresponding CV for each type of neuron at each modulatory condition (color coded). **(E)** Average spike frequency (*F*_spks_) within a burst and corresponding CV for each type of neuron at each modulatory condition (color coded). N = 25 animals. Individual dots represent data from individual experiments, red bars indicate the (circular) mean. Asterisks indicate significant differences between groups (Dunn’s or Tukey’s post-hoc test (see Methods), p ≤ 0.05). Dagger indicates that this group is different from all others, except other groups marked with dagger. Total modulator concentration for [mid]: 3 x 10^−8^ M. ANOVA results in Table 3. Figure 5-1: Raw data for all panels.

**Table 3:** ANOVA results for pyloric rhythm attributes mid modulator concentrations (different dataset than in Table 2). If tests for normality or equal variance failed, results are for ANOVA on ranks. Groups are intact, decentralized, P [mid], PC [mid], PCR [mid], wash. p values smaller than α are printed in bold. N: number of animals; df: degrees of freedom; res: residual.

| N | attribute | normality | variance | test statistic | df | res. | p |
| --- | --- | --- | --- | --- | --- | --- | --- |
| 25 | Cycle frq | fail |  | H = 61.109 | 5 |  | **<0.001** |
| 25 | PD off | fail |  | H = 33.152 | 5 |  | **<0.001** |
| 25 | LP on | pass | fail | H = 37.630 | 5 |  | **<0.001** |
| 25 | LP off | pass | fail | H = 61.122 | 5 |  | **<0.001** |
| 25 | PY on | pass | pass | F = 7.677 | 5 | 134 | **<0.001** |
| 25 | PY off | pass | pass | F = 7.224 | 5 | 134 | **<0.001** |
| 25 | PD spikes | fail |  | H = 31.084 | 5 |  | **<0.001** |
| 25 | PD spike frq | fail |  | H = 5.576 | 5 |  | 0.350 |
| 25 | LP spikes | pass | fail | H = 94.633 | 5 |  | <0.001 |
| 25 | LP spike frq | pass | pass | F = 29.448 | 5 | 131 | <0.001 |
| 25 | PY spikes | fail |  | H = 41.5592 | 5 |  | <0.001 |
| 25 | PY spike frq | pass | pass | F = 3.009 | 5 | 134 | 0.013 |

### 6.2 Compared to single peptide modulators, comodulation at mid concentrations did not reduce interindividual variability at the single-neuron level

In a previous study we showed that neuropeptide modulation at saturating concentration reduces the interindividual variability of excitability attributes of the synaptically isolated LP neuron (Schneider et al. 2022). This finding raises the possibility that the reduction of circuit output variability arising by increasing numbers of comodulatory peptides at mid-level concentration may be a consequence of a similar reduction at the level of individual pyloric neurons. To examine this hypothesis, we measured single-neuron interindividual variability using the same protocols as in Schneider et al. (2022), but with increasing numbers of comodulators applied at mid concentration. We focused on the synaptically isolated LP neuron because it exists only in a single copy in each animal, and it is targeted by all three neuropeptides that we used in this study (Swensen and Marder 2001).

The first excitability attribute that we examined was the spike frequency vs. input current relationship (*f-I* curve). We measured the *f-I* curves by applying increasing and decreasing current steps and measured the firing frequency at each step (Fig. 6A). In order to compare these data across individuals, we parameterized the *f-I* curves across the range of applied currents by fitting these data with a sublinear power function *f* (*I*) = *a*(*I* − *I*)*^b^* where *I*_0_ is the zero-intercept, *a* is the slope and the power *b* represents the curvature between 0 and 1, ensuring the curve is at most linear (Fig. 6B fit curve; also see Methods Eq. 1). Both the slope factor *a*, and the exponent *b*, but not *I_0_*, were different in mean value in the presence of neuropeptides compared to the unmodulated condition, indicating a higher LP spike rate in the presence of the neuropeptides (ANOVA results in Table 4; raw data are provided in Figure 6-1). In general, fit parameter variability was lower when one or more peptide modulators were present, but there was no consistent reduction in variability with the addition of comodulators (Fig. 6C).

**Figure 6:**
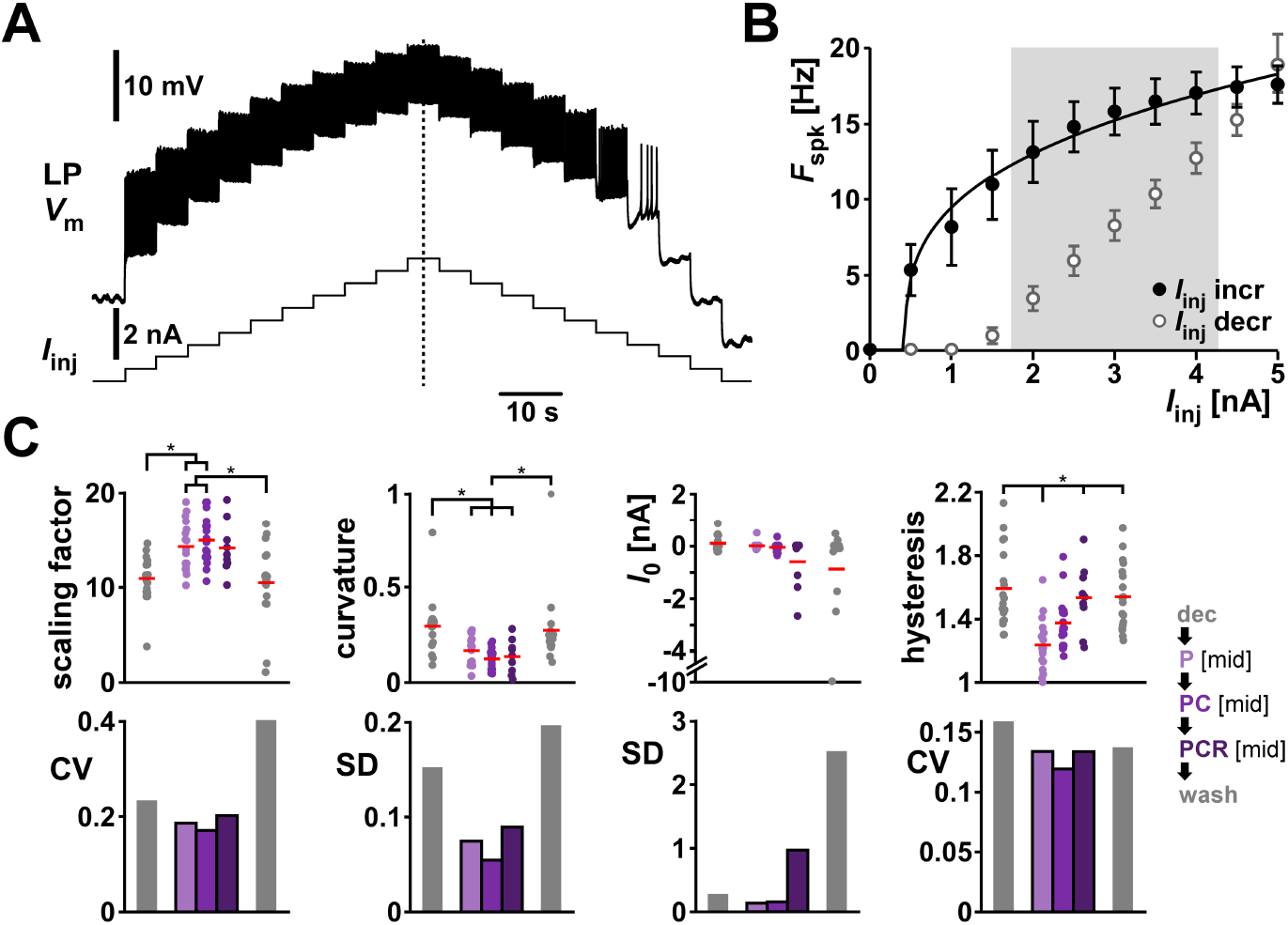
Comodulation does not reduce the interindividual variability of excitability metrics on the single cell level. **(A)** Voltage changes of one example LP neuron in response to increasing and decreasing current injection. **(B)** *f-I* curve of the experiment in (A). The average spike frequency at each current level (dots and error bars indicate mean ± SD) was fitted with a power function (black line). To calculate *f-I* hysteresis between increasing (filled circles) and decreasing (open circles) levels of current injection, we divided the average spiking frequency in increasing by decreasing current levels between 2 and 4 nA (shaded area). **(C)** Fit parameters (scaling factor = *a*, curvature = *b* in Eq. 1). and hysteresis with the corresponding metric of variability (CV, or SD for interval data) for different modulatory conditions (color coded). Dots represent values from individual experiments, horizontal bars indicate the mean value. N = 19, except PCR N = 11. Asterisks indicate significant differences between groups (Dunn’s post-hoc test, p ≤ 0.05). Total modulator concentration for [mid]: 1-3 x 10^−8^ M for P and PC, 3 x 10^−8^ M for PCR. ANOVA results in Table 4. Figure 6-1: Raw data for panel C.

**Table 4:** ANOVA (on ranks) results for f-I parameters. Groups are decentralized, P [mid], PC [mid], PCR [mid], wash. p values smaller than α are printed in bold. N: number of animals; df: degrees of freedom.

| N | parameter | normality | test statistic | df | p |
| --- | --- | --- | --- | --- | --- |
| 17 | a | fail | H = 24.667 | 4 | **<0.001** |
| 17 | b | fail | H = 29.801 | 4 | **<0.001** |
| 17 | $I_0$ | fail | H = 3.680 | 4 | 0.451 |
| 19 | hysteresis | fail | H = 27.796 | 4 | **<0.001** |

As seen in the voltage trace of Figure 6A, the *f-I* curve shows hysteresis, in that the firing frequency for each current step depends on whether it is measured on the increasing or decreasing leg of injected currents (Fig. 6B). We quantified hysteresis, as detailed in the Methods, by calculating the ratio between the mean *f* value on the increasing and decreasing legs of injected currents within the shaded region (2 –4 nA) of Fig. 6B. Thus, a value greater than 1 means that LP was spiking at a higher frequency when the current changed in increasing direction. Even though neuropeptides influenced hysteresis, they did not change its variability (Fig. 6C).

LP is a follower neuron and, during normal pyloric activity, generates bursts of spikes upon rebound from inhibition. To examine the post-inhibitory rebound properties of LP, we hyperpolarized the isolated LP neuron for 10 s and characterized the spiking activity in the 10 s window after release from inhibition in different modulatory conditions (Fig. 7A). We did so by measuring the latency from the end of inhibition to the first spike, and generated spike histograms from the 10 s window following the inhibition (Fig. 7B). We characterized the spike histogram by fitting a sigmoid function to the cumulative spike count (Fig. 7C).

**Figure 7:**
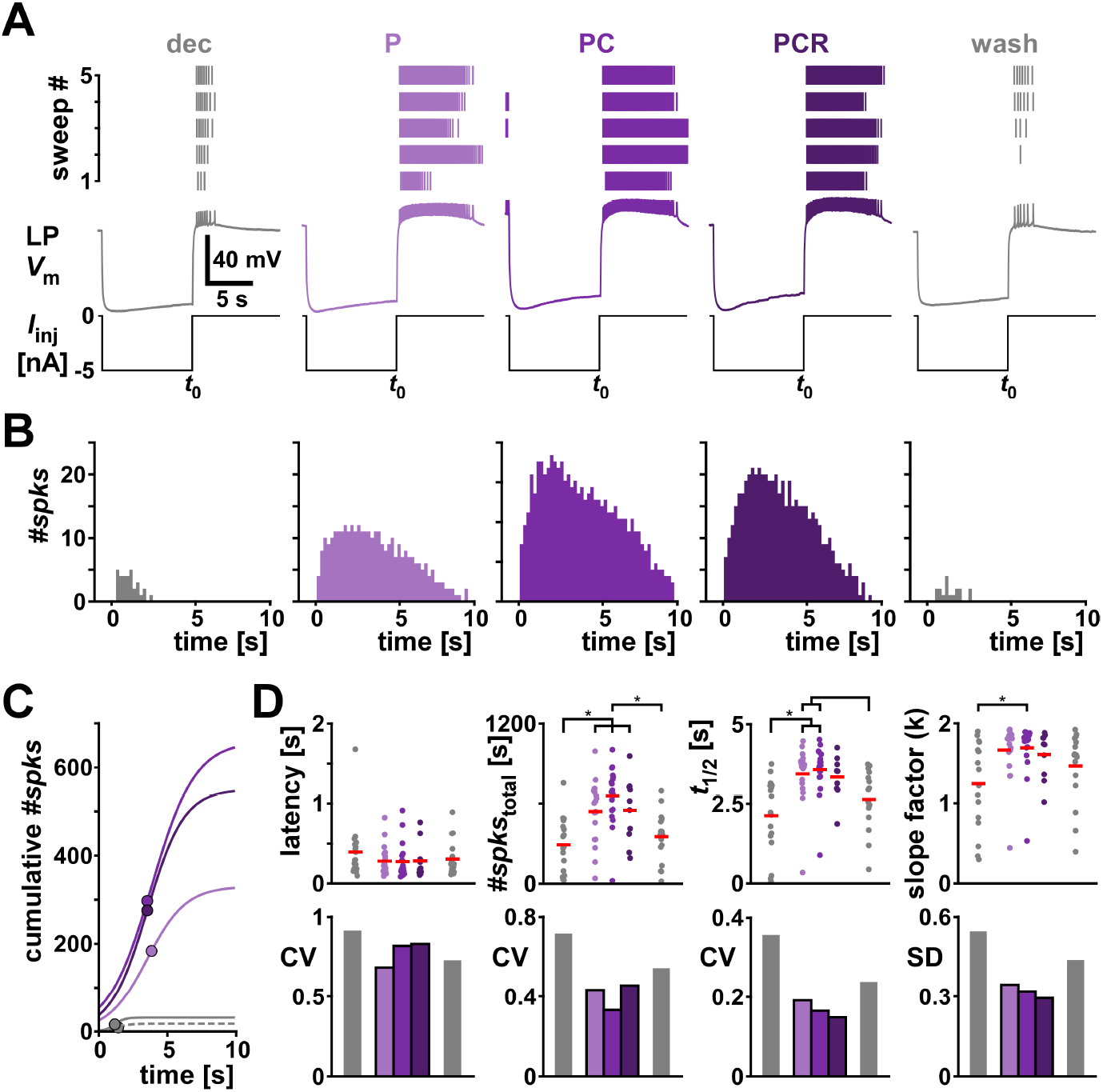
Comodulation does not add to the reduction of interindividual variability of rebound metrics on the single cell level. **(A)** An example of the rebound experiment in different modulatory conditions (color coded). Top panels show the spike raster plots for all 5 sweeps. Bottom shows the intracellular voltage of one example sweep. t_0_ indicates the time point when LP was released from current injection. **(B)** Spike histograms across all sweeps in different modulatory conditions (color coded). **(C)** Sigmoid fits to the cumulative spike histograms. Dots indicate the sigmoid midpoint. Dashed grey line indicates wash. **(D)** Latency and fit parameters (#*spikes*_total_ = a, slope factor = *k* in Eq. 2) with the corresponding metric of variability (CV, or SD for interval data) for different modulatory conditions (color coded). Dots represent values from individual experiments, horizontal bars indicate the mean value. N = 19, except PCR N = 11. Asterisks indicate significant differences between groups (Dunn’s or Tukey’s post-hoc test (see Methods), p ≤ 0.05). Total modulator concentration for [mid]: 1-3 x 10^−8^ M for P and PC, 3 x 10^−8^ M for PCR. ANOVA results in Table 5. Figure 7-1: Raw data for panel D.

**Table 5:** ANOVA results for 10 s rebound parameters. If tests for normality or equal variance failed, results are for ANOVA on ranks. Groups are decentralized, P [mid], PC [mid], PCR [mid], wash. p values smaller than α are printed in bold. N: number of animals; df: degrees of freedom; res: residual.

| N | parameter | normality | variance | test statistic | df | res. | p |
| --- | --- | --- | --- | --- | --- | --- | --- |
| 19 | latency | fail |  | H = 3.385 | 4 |  | 0.495 |
| 19 | a | pass | pass | F = 8.665 | 4 | 77 | **<0.001** |
| 19 | $t_{1/2}$ | fail | | H = 28.343 | 4 | | **<0.001** |
| 19 | k | fail |  | H = 13.148 | 4 |  | 0.011 |

Adding one or several neuropeptides at mid concentration significantly changed the mean values of the fit parameters of the sigmoid but not the rebound latency (Fig. 7D top row; ANOVA results in Table 5; raw data are provided in Figure 7-1). All fit parameters increased with neuropeptides compared to the decentralized control condition, indicating that LP was overall firing more spikes in longer bursts, consistent with the increased excitability seen in the *f-I* curves. Neuropeptides increased the likelihood of LP rebound from inhibition and overall decreased the variability of the rebound parameters compared to the two unmodulated conditions (decentralized and wash). However, we observed a decrease of variability with increasing numbers of neuropeptides only for the fit parameters *t_1/2_* and *k*, but not for the latency or the total number of spikes within a rebound burst (Fig. 7D bottom row). The decrease of variability was present even when the parameters were not statistically different between the modulatory conditions.

In the intact pyloric circuit, LP is periodically inhibited by a group of pacemaker neurons and typical values of pyloric *F*_cycle_ range from 0.5 to 2 Hz. To mimic this effect, we modified the rebound protocol and hyperpolarized for 1 s every 2 s, repeated 20 times (Fig. 8A). As shown previously (Anwar et al. 2022; Goaillard et al. 2010; Schneider et al. 2022), LP rebound only reaches steady state after several cycles (Fig. 8B). Therefore, we only included the last 10 sweeps in our analysis and examined the same rebound parameters as before. (Fig. 8C). In contrast to the 10 s rebound protocol, the application of neuromodulators at mid concentration did not significantly change any of the parameters with this faster protocol (Fig. 8D, ANOVA results in Table 6; raw data are provided in Figure 8-1). Furthermore, while the metrics of variability did not increase with the addition of comodulators, the reduction of variability across animals was minor and did not recover after washing (Fig. 8E).

**Figure 8:**
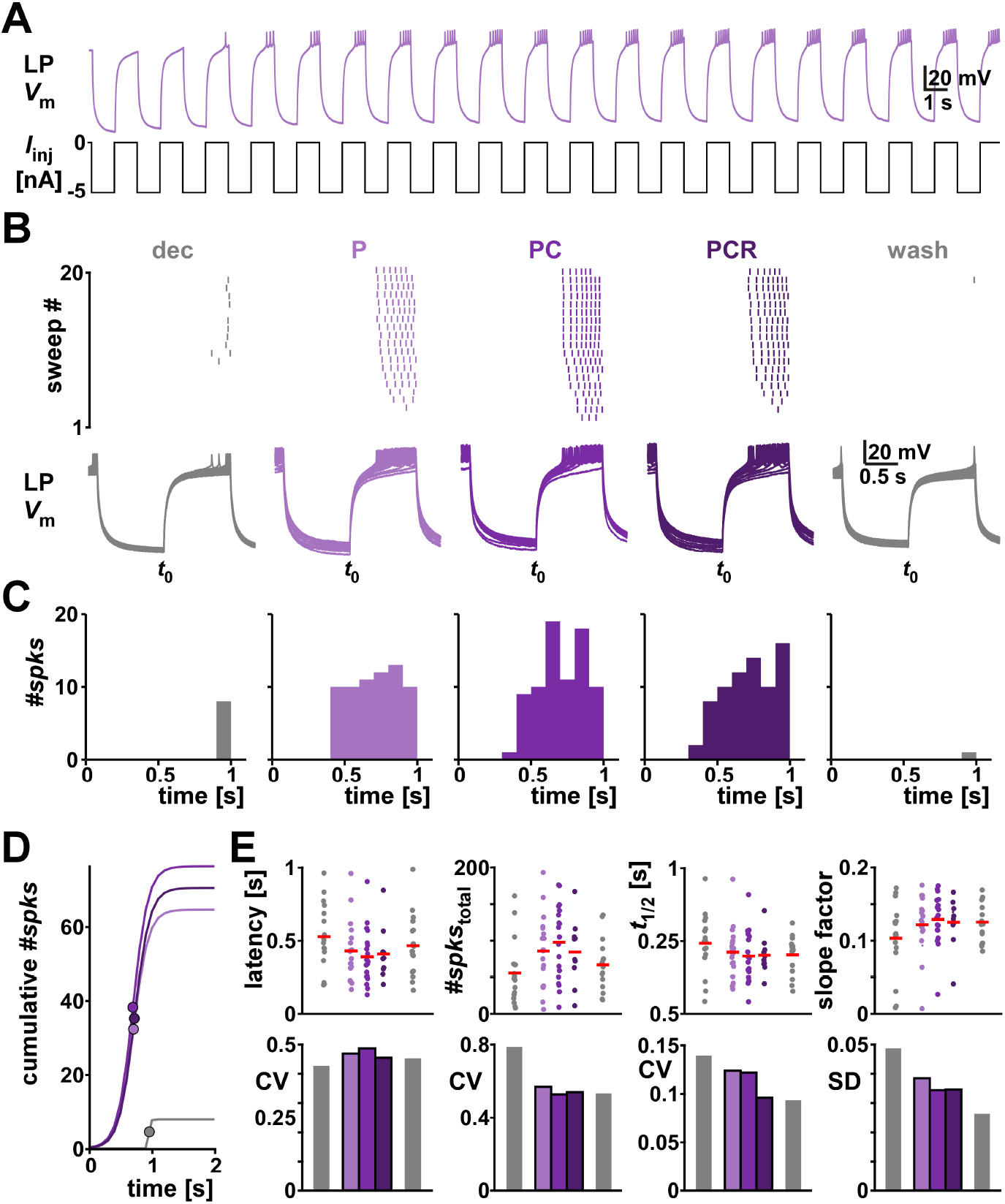
Comodulation does not reduce the interindividual variability of rebound from periodic inhibition on the single cell level. **(A)** One example of LP membrane potential in response to periodic hyperpolarization. **(B)** One example periodic rebound experiment in different modulatory conditions (color coded). Top panels show the spike raster plots for all 20 sweeps. Bottom shows the intracellular voltage of all sweeps. t_0_ indicates the time point when LP was released from current injection. **(C)** Spike histogram across the last 10 sweeps in different modulatory conditions. **(D)** Sigmoid fits to the cumulative spike histograms. Dots indicate the sigmoid midpoint. **(E)** Latency and fit parameters (#*spikes*_total_ = a, slope factor = *k* in Eq. 2) with the corresponding metric of variability (CV, or SD for interval data) for different modulatory conditions (color coded). Dots represent values from individual experiments, horizontal bars indicate the mean. N = 19, except PCR N = 11. No significant differences between groups (ANOVA results in Table 6.). Total modulator concentration for [mid]: 1-3 x 10^−8^ M for P and PC, 3 x 10^−8^ M for PCR. Figure 8-1: Raw data for panel E.

**Table 6:** ANOVA results for 1 s rebound parameters. If tests for normality or equal variance failed, results are for ANOVA on ranks. Groups are decentralized, P [mid], PC [mid], PCR [mid], wash. p values smaller than α are printed in bold. N: number of animals; df: degrees of freedom; res: residual.

| N | parameter | normality | variance | test statistic | df | res. | p |
| --- | --- | --- | --- | --- | --- | --- | --- |
| 19 | latency | fail |  | H = 5.329 | 4 |  | 0.249 |
| 19 | a | pass | pass | F = 2.371 | 4 | 76 | 0.060 |
| 19 | $t_{1/2}$ | pass | pass | F = 0.765 | 4 | 76 | 0.551 |
| 19 | k | fail |  | H = 4.068 | 4 |  | 0.397 |

### 6.3 Peptide comodulation at saturating concentrations did not reduce variability of circuit activity attributes beyond the effects of a single peptide

So far, we have shown in this study that increasing the number of comodulatory neuropeptides at mid concentrations, but not at low concentrations, decreased the variability of several pyloric circuit rhythm attributes across animals. In contrast, at the single neuron level, additional mid-concentration comodulatory reduction of variability is absent or ambiguous even though peptide modulation does reduce variability. This latter finding is consistent with our previous findings that peptide modulation at saturating concentrations (at or above 1 µM) reduces single-neuron variability but addition of a second comodulatory peptide does not further reduce it (Schneider et al. 2022). However, at saturating concentrations, convergent modulators may also occlude one another’s actions. This raised the question of whether comodulation reduces variability of the full circuit output attributes.

We addressed this question with the same two modulators used in the Schneider et al (2022) study. Figure 9 summarizes the results of applying P and PC at a total concentration of 1 µM to the decentralized preparation (ANOVA results in Table 7; raw data are provided in Figure 9-1). In contrast to the experiments at low and mid concentrations, we only used experiments where the pyloric rhythm continued when the preparation was decentralized. In these experiments, *F*_cycle_ was significantly higher in the intact condition (Fig. 9A), and decentralization impacted the off phases of PD and LP and the on phase of PY (Fig. 9B). For frequency and phases, variability across individuals was lower with modulators than without (Fig. 9A-B). Additionally, the variability at high concentrations was even lower than in three modulators at mid concentration (indicated by the hollow purple bars in Fig. 9A-C bottom row). This was also the case for the sum of the eigenvalues of the phase covariance matrix (Fig. 9C). At saturating concentrations, the variability across animals was in the same range as in the intact condition. But even though the overall variability of frequency and phase was lower at high concentrations, the addition of a comodulator did not further reduce variability in a consistent manner as it did at mid concentrations. Variability metrics for spike number (Fig. 9D) and spike frequency (Fig. 9E) within a burst, however, decreased with the addition of a comodulator but the variability in high modulator concentrations was comparable to that in mid concentrations. Taken together, these results show that comodulation at saturating concentrations is not as consistently reducing variability at the network level as comodulation below saturating concentrations.

**Figure 9:**
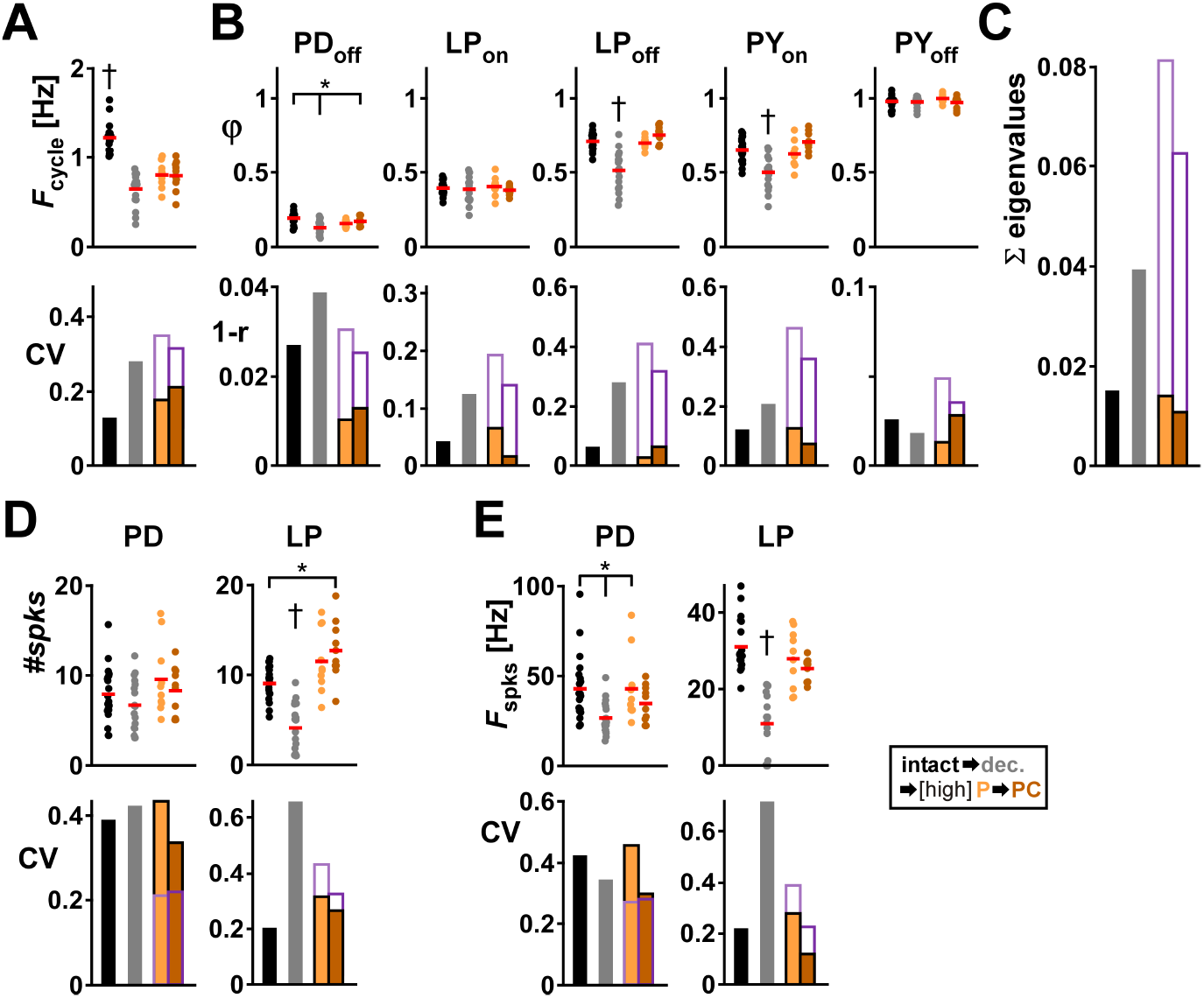
Neuropeptides at high concentration reduce the interindividual variability of pyloric rhythm parameters on the circuit output level. **(A)** *F*_cycle_ and the corresponding CV under different modulatory conditions (color coded). **(B)** Burst start (on) and termination (off) and the corresponding circular variance (1-r, see Methods) under different modulatory conditions (color coded). **(C)** Sum of eigenvalues from the covariance matrix of the phases for each modulatory condition. **(D)** Number of spikes per burst and corresponding CV for each type of neuron at different high concentration modulatory conditions (color coded). **(E)** Average Spike frequency within a burst and corresponding CV for each type of neuron at each modulatory condition (color coded). Separate datasets for P and PC, N = 10 each. Individual dots represent data from individual experiments, red bars indicate the mean value. Asterisks indicate significant differences between groups (Dunn’s or Tukey’s post-hoc test (see Methods), p ≤ 0.05). Dagger indicates that this group is different from all others. The purple empty bars indicate the value of the variability metric at [mid] PCR from Figs. 2 to 4. Total modulator concentration for [high]: 10^−6^ M. ANOVA results in Table 7. Figure 9-1: Raw data for all panels.

**Table 7:** ANOVA results for pyloric rhythm attributes at high modulator concentrations. If tests for normality or equal variance failed, results are for ANOVA on ranks. Groups are intact, decentralized, P [high], PC [high], wash. p values smaller than α are printed in bold. N: number of animals; df: degrees of freedom; res: residual.

| N | attribute | normality | variance | test statistic | df | res. | p |
| --- | --- | --- | --- | --- | --- | --- | --- |
| 20 | Cycle frq | pass | pass | F = 43.965 | 3 | 54 | **<0.001** |
| 20 | PD off | pass | pass | F = 9.373 | 3 | 54 | **<0.001** |
| 20 | LP on | pass | fail | H = 1.457 | 3 |  | 0.692 |
| 20 | LP off | pass | fail | H = 30.555 | 3 |  | **<0.001** |
| 20 | PY on | pass | pass | F = 14.001 | 3 | 54 | **<0.001** |
| 20 | PY off | fail |  | H = 2.913 | 3 |  | 0.405 |
| 20 | PD spikes | fail |  | H = 4.285 | 3 |  | 0.232 |
| 20 | PD spike frq | fail |  | H = 13.449 | 3 |  | **0.004** |
| 20 | LP spikes | pass | pass | F = 28.269 | 3 | 54 | **<0.001** |
| 20 | LP spike frq | pass | pass | F = 30.227 | 3 | 54 | **<0.001** |

### 6.4 Does excitatory comodulation generically reduce variability?

As stated above, excitatory modulatory peptides influence the pyloric circuit by activating the voltage-gated inward current, *I*_MI_, and by increasing synaptic efficacy. Because these peptide modulators have distinct but overlapping targets, the effect of additive comodulation would be to primarily activate *I*_MI_ in additional pyloric neurons, and to modulate more synapses. Increasing the levels of *I*_MI_ in a neuron would increase its excitability. Is it possible then that simply targeting more pyloric neurons with higher and higher levels of *I*_MI_ would generically (i.e., without additional constraints on the system) result in a reduction of interindividual variability? We examined this possibility using computational modeling.

It is well established that the basic triphasic pyloric pattern can be generated in models with widely varying levels of ion current maximal conductances (Prinz et al. 2004). We therefore started with a family of 175 pyloric model circuits, each with the same neuron types and connectivity (Fig. 1B). These neurons expressed the same cohort of ionic currents (Fig. 10A, B), but with a distinct combination of maximal conductance values (see Methods for a detailed description; equations and parameters in Figures 10-1, 10-2, and 10-3). We modeled peptide neuromodulation by increasing the maximal *g*_MI_ levels in each target neuron (Fig. 10B). We ensured that each model circuit expressed a triphasic motor pattern similar to that observed in the biological circuit (Fig. 10C). Variability of circuit output attributes was measured as the circular variance (for phases, as in the biological experiments) or CV (for all other attributes) across the 175 model circuits, under different levels of modulatory input.

**Figure 10:**
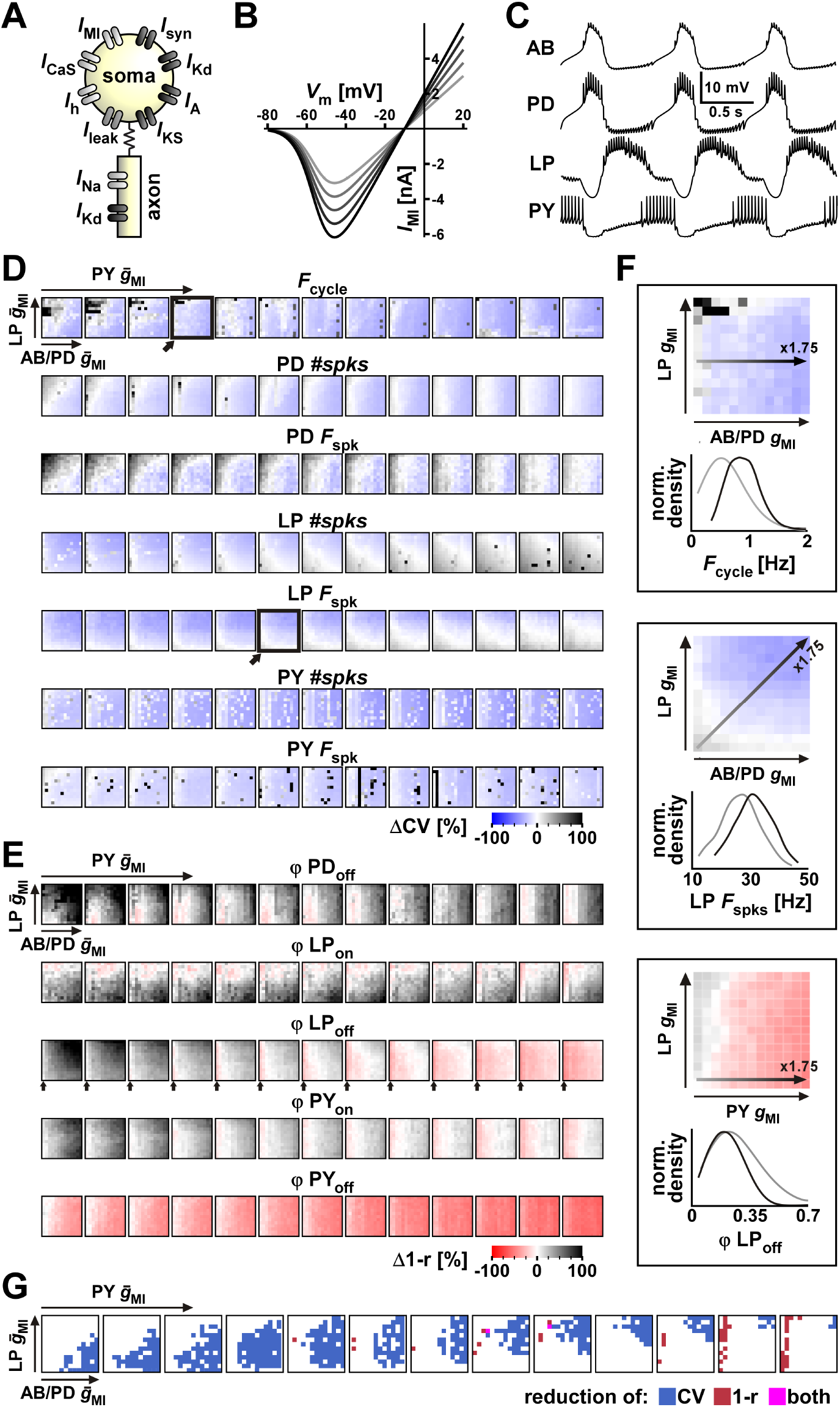
Changes in variability of burst attributes and activity phases depend on the regulation of *I*_MI_ levels across different neurons in a family of model circuits. **(A)** Schematic diagram of neurons and ionic currents included in the model. **(B)** The modulatory current *I*_MI_ *I-V* relationship as modeled. Different levels of modulatory input change the maximal conductance *g*_MI_, resulting in the changes in the *I-V* relationship as shown (increase shown from grey to black). **(C)** Example voltage activity of the four neuron types used in the models at a mid-level modulation of *g*_MI_ for all neurons shows the triphasic pyloric activity. **(D)** Change in the variability (measured as CV) of the circuit output attributes, *F*_cycle_, LP #*spks*, LP *F*_spks_, PY #*spk*, PY *F*_spks_ across the 175 model circuits, under different levels of modulatory input. Modulatory input is modeled by increasing *g*_MI_ from its baseline unmodulated value, from 1x to 1.75x in 13 multiplicative steps, across the whole family of model circuits. *g*_MI_ was increased independently in each of the three neuron groups (pacemakers AB/PD, LP, PYs), resulting in a 3D grid. For each attribute (row) each panel represents the increase in pacemaker *g*_MI_ on the horizontal axis and the increase in LP *g*_MI_ on the vertical axis. The increase in PY *g*_MI_ is represented in different panels from left to right. Change in CV (with respect to the unmodulated state) is plotted as a heatmap where colors from blue-to-white-to-black represent decreased-no change-increased CV (see scalebar) of that attribute. Therefore, the left bottom square (white) of the leftmost panel shows the fully unmodulated family of models and the top right square of the rightmost panel shows the fully modulated family. **(E)** Change in the variability (measured as circular variance 1-*r*) of the burst phases under different levels of modulatory input. Description is as in panel D, but with red replacing blue for reduction in variability. **(F)** Zoom-in to three representative panels of the 3D reduction of variability grid. Data shown are for three attributes: *F*_cycle_ (top box, corresponding to arrow-highlighted panel in the top row of D), LP *F*_spks_ (middle box, corresponding to arrow-highlighted panel in the third row of D) and ϕLP_off_ (bottom box, corresponding to columns indicated by arrows in each panel in the third row of E). In each group, the change in the normalized distribution of that attribute is shown when *g*_MI_ is changed along the path and direction shown by the arrow in the colored grid panel. This highlights how the density function narrows (representing reduction in variability) by increasing *g*_MI_. **(G)** Regions of reduction in variability for all circuit output attributes. Data are represented as in each row of D or E. Each blue dot represents a point in modulatory parameter space where the variability of all attributes in panel D is reduced by at least 2.5%. Each red dot represents the reduction of variability of all phases in panel E. Each pink dot shows the overlap between blue and red, indicating that reduction of variability of all circuit output attributes occurs only sporadically. Figure 10-1: Mechanistic equations describing the model family. Figure 10-2: Parameters for intrinsic ionic current mechanisms of the model equations. Figure 10-3: Parameters for synaptic mechanisms of the model equations.

To characterize the effects of comodulation, we assumed first that each modulator targets only one of three groups of neurons. We did so by splitting the circuit neurons into the following three groups: the pacemaker (AB and PD) neurons, the LP neuron and the PY neurons. We started with the family of model circuits in the unmodulated state and then increased *g*_MI_ independently in each of the three groups. For each group, *g*_MI_ was increased from its baseline unmodulated value, in multiplicative steps, across the whole family of model circuits. Because *g*_MI_ was increased independently in each of the three groups, we obtained a 3D grid representation of our modulatory space (Fig. 10C). Note that the effect of any neuromodulator that targets more than one group by activating different levels of *I*_MI_ in different neuron groups would also be represented in this grid.

We analyzed the effect of neuromodulators on variability for each attribute by plotting how variability was reduced or increased across the 3D grid (Fig. 10D, E). The variability of *F*_cycle_ was reduced by increasing *g*_MI_ in the pacemakers (as seen in the increase in the blue color along the horizontal axis of each box of the top row in Figure 10D). Increasing *g*_MI_ in LP, however, increased the variability of *F*_cycle_, but only when *g*_MI_ values were low in PY (as seen in the increase in the black color along the vertical axis of the leftmost 4 boxes of the top row in Figure 10D). The variability of LP spike number and spike frequency was insensitive to *g*_MI_ in the pacemakers, reduced with *g*_MI_ in LP and increased with *g*_MI_ in PY. The variability of PY spike number and frequency decreased with *g*_MI_ in the pacemakers but was mostly insensitive to *g*_MI_ in LP or even in the PYs themselves. This is probably because PY spiking variability was most sensitive to the variability of *F*_cycle_.

The variability of the burst onset (ϕ_on_) and end (ϕ_off_) phases showed a remarkably different pattern than that of *F*_cycle_ or neuronal spiking activities (Fig. 10E). Surprisingly, ϕPD_off_ (i.e., the PD duty cycle) became more variable with increased *g*_MI_ in the pacemakers. Its variability also increased with *g*_MI_ in LP but was reduced by *g*_MI_ in PY. ϕLP_on_ became less variable with *g*_MI_ in LP and to a smaller extent with *g*_MI_ in PY, but increased in variability with *g*_MI_ in the pacemakers, probably because it is closely tied to ϕPD_off_. In contrast, ϕLP_off_ variability increased with *g*_MI_ in the pacemakers, but this was compensated for by *g*_MI_ in PY which greatly reduced this variability. Somewhat surprisingly, ϕLP_off_ variability was insensitive to *g*_MI_ in LP. ϕPY_on_ became more variable with *g*_MI_ in either the pacemakers or LP, but this was compensated for by *g*_MI_ in PY itself when the pacemaker *g*_MI_ was not too strong. In our models, ϕPY_off_ was very close to 1 (the onset of the next cycle) and did not show much variability.

To understand how increasing *g*_MI_ leads to reduction of variability, we examined how the distribution of the values of an output attribute may change as a function of *g*_MI_. We show three examples of this effect in Figure 10F. Increasing *g*_MI_ in the pacemakers shifted the distribution (shown as normalized probability density function) of *F*_cycle_ to larger values, but across the family of models, *F*_cycle_ remained below an upper bound of ∼2 Hz (Fig. 10F top panels). Thus, the rising segment of the density function curve shifted to the right much more than its falling segment, resulting in a narrower density function and therefore a lower CV. A similar narrowing of the density function occurred for LP spike frequency, when *g*_MI_ was simultaneously increased in the pacemakers and LP (Fig. 10F middle panels). For the activity phases, reduction of variability could also result from shifting the density function to the left. For example, increasing *g*_MI_ in PY could result in restricting the maximum value of ϕLP_off_ without changing its minimum (Fig. 10F bottom panels). This occurred because the end of the LP burst is in great part due to the inhibition it receives from the PY neurons (see Fig. 10C) and strengthening the PY burst (earlier burst onset and more spikes per burst) by increasing *g*_MI_ would constrain the end of the LP burst.

To see how comodulation of the model circuits resulted in an overall reduction of variability of circuit activity, we plotted the overlapping regions in which variability of individual output attributes was reduced (Fig. 10G). We did this separately for activity phases (circular variances, shown in red) and all other attributes (CVs, shown in blue). The resulting graph provided two sets of information. First, it showed that as levels of *g*_MI_ increase across all three neuron groups, akin to increased comodulation, there is reduction of variability of all CVs but only up to a certain *g*_MI_ level, akin to an intermediate concentration (increase in blue regions from bottom left to top right in each panel and from left to right across panels). Second, it showed that although increasing *g*_MI_ in all three neuron groups could also reduce the circular variance of all phase attributes (red), the range of *g*_MI_ for which this overall reduction occurred had little overlap with the region in which CVs of other attributes were reduced (overlap shown in magenta). We therefore conclude that the reduction of variability across both phase and other attributes that we observed in the biological circuit do not arise generically with our simplifying assumptions used to model the circuit and its modulation.

## 7 Discussion

### 7.1 How could convergent comodulation reduce interindividual variability?

Generally, there are two possibilities for how convergent peptide comodulation of the pyloric circuit could lead to reduction of circuit output variability. First, variability may be reduced at the single-neuron level, for instance because comodulation produces more consistent levels of *I*_MI_, as some neuron types only have receptors for a subset of these peptides (Garcia et al. 2015; Swensen and Marder 2001). Alternatively, reduction in interindividual variability could be a circuit-level phenomenon.

Recently, we examined the interindividual variability of single neuron responses to excitatory neuromodulation (Schneider et al. 2022). We used the LP neuron whose natural oscillations in the intact system show remarkable similarity across animals (Schulz et al. 2006). When synaptically isolated, LP neuron responses to stimuli were quite variable across preparations, but this variability was greatly reduced in the presence of PROC, which activates *I*_MI_ in LP and increases its overall excitability (Schneider et al. 2021). A simple modeling analysis showed that increasing excitability in a family of such neurons restricts the *f-I* relationship both by increasing the minimum bound on the firing rate, and contracting the range of firing rates values at a given current level. Although the *f-I* relationship was one of many neural output attributes we examined, the effect of an excitatory neuromodulator on this relationship hinted at a more general principle through which increase in excitability of a neuron could constrain variability of its output attributes. The statistical distribution of any biological output attribute (frequency, phase, etc.) across individuals is constrained by a lower and an upper limit. A neuromodulator may influence the shape of this distribution by shifting these limits. If such a shift brings these limits closer together without substantially changing the mean value, it could lead to a reduction in variability (variance or CV). This informal argument also shows how understanding the constraints on variability can be moved from a statistical viewpoint to a mechanistic explanation in which modulator actions on ionic mechanisms, together with biophysical constraints, can unmask the modulator effects on variability.

In the same study, the reduction of variability was the same in the presence of PROC alone or with the combined application of PROC and CCAP, indicating that comodulation did not reduce variability compared to a single modulator. In the current study, we found that peptide modulation at mid-level concentrations of 30 nM also reduced variability of the isolated LP neuron outputs compared to the unmodulated states and there was no additional reduction of variability by comodulation (Figs. 6-8). Convergent modulator effects on a neuron depend on concentration and receptor expression. If two modulators acting on the same downstream pathway are applied at saturating concentrations, occlusion occurs and comodulation will have little additional effect compared to single modulator application. However, additive application will increase excitability at sub-saturation levels, which is why we used constant total concentrations. If the activation pathway towards the common target ion channel is through multiple different receptor populations, interindividual variability may be decreased by comodulation because variability in the different receptor populations averages out.

Yet, increasing the number of modulators while keeping the total concentration constant is an imperfect way to keep the overall activation of a circuit similar for several reasons. First, receptor activation depends on a dose-response curve that is typically sigmoidal on a log scale. Therefore, even for two receptors with identical expression levels and concentration-dependence, application of a single modulator at any concentration results in a lower level of overall G protein activation than co-application of two modulators at half that concentration. Second, receptor expression and dose-response curves vary across neuron types (Garcia et al. 2015), rendering consequences of co-applications further distinct from simple linearly additive effects. An extreme version of this is when a single neuron only expresses the receptor to one of two co-applied modulators, resulting in less activation during co-application than during single application. Third, co-application of neuropeptides may lead to sublinear combined *I*_MI_ activation because of potential inhibitory interactions in the signaling pathways (Li et al. 2018). For these reasons, we do not claim that keeping the total concentrations constant ensures equal overall circuit activation. However, it is a practical way to keep the circuit in an activation range that is not subject to saturation effects, which would more easily occur with additive applications.

Still, comodulation can lead to increasingly consistent modulation across multiple components at the circuit level, both with additive and constant total concentrations. Both increase in excitability and consistency in activation can decrease interindividual variability. Thus, our findings indicate that the comodulation-mediated reduction of variability is unlikely to emerge at the level of individual neurons and therefore it likely emerges at the circuit level.

There are many ways such circuit-level actions of modulators can arise. For instance, modulation of a single neuron may have effects that are restricted to that neuron or, conversely, reverberate through other circuit neurons, as was demonstrated in a computational study exploring the effects of hub neurons in a network (Gutierrez and Marder 2014). Additionally, the excitatory neuropeptides also influence synaptic efficacy and dynamics in this circuit (Atamturktur and Nadim 2011; Li et al. 2018; Thirumalai et al. 2006; Zhao et al. 2011), which in turn influences the activities of all circuit neurons. Finally, interactions between nonlinear properties of bursting neurons connected with recurrent synapses that have short-term dynamics could give rise to effects that are not present in isolated neurons or feedforward circuits (Akcay et al. 2018; Tsodyks et al. 2000; Yuste 2015).

### 7.2 What the pyloric circuit models say about the reduction in interindividual variability

The purpose of our modeling exercise in this study was to see if convergent comodulation would generically lead to reduction of variability. We did not model the peptide targets as in the biological circuit (i.e., as shown in Fig. 2A), but as a grid-based increase of *g*_MI_ in individual circuit neurons. Yet this grid-based model version of progressive comodulation includes, as a subset, the biological cases in which an increasing number of circuit neurons were targeted by one, two or three convergent modulators (as in Fig. 2B).

This modeling exercise led to several results. First, it showed that increasing *g*_MI_ levels in an increasing number of neurons can in fact lead to a reduction of output attribute variability (Fig. 10D, E). Interestingly, examining the effect of comodulation on the normalized probability distribution of some circuit output attributes confirmed the intuition that reduction of variability could simply result from constricting the upper or lower limits of these distributions (Fig. 10F).

Second, the modeling confirmed that, *a priori*, there is no reason that reduction of variability of different circuit output attributes should overlap. The way that increasing levels of *g*_MI_ in different neurons reduces variability of one output attribute (say, *F*_cycle_) is not the same way that increasing *g*_MI_ would decrease variability in another attribute (say, ϕLP_on_). In fact, even using the limited condition that variability of all attributes be reduced by only 2.5% resulted in a very small number of modulator combinations that satisfied an overall reduction of variability of several attributes. Additionally, there was almost no overlap between the combination of *g*_MI_ levels (i.e., comodulation of) all neurons that resulted in a simultaneous reduction of variability in *F*_cycle_ and spiking activity on the one hand and burst phases on the other (Fig. 10G). Thus, the overall lesson of the model is that simply setting the connectivity and tuning a modestly biophysically realistic model to produce activity within the biological range is not sufficient to explain how progressively increasing comodulation results in reduction of variability in almost all circuit output attributes as we see in our biological data. This may mean that our family of models did not include some important aspects of the modulatory actions, such as the effects of synaptic efficacy, or that some unknown additional mechanisms in the biological circuit allow for such a robust reduction of variability. Both these possibilities could be explored by a more expansive computational modeling analysis aided by more focused smaller scale mathematical models to elucidate the mechanisms that may shape the distributions of different output attributes.

### 7.3 Modulator action to shape specific circuit outputs vs. reduction of variability

Neuromodulators play a crucial role in the flexibility of neural circuits. This diverse set of chemicals act on a wide range of (mainly metabotropic) receptors to alter levels of leak and voltage-gated ionic currents and even ion transporters, thereby greatly influencing the intrinsic excitability of neurons (Marder 2012). The same receptors also act on synaptic release mechanisms and neurotransmitter receptors to enhance or suppress synaptic transmission, alter the sensitivity of neurons to input, and modify synaptic plasticity (Nadim and Bucher 2014). This dynamic modulation allows neural circuits to adapt and reorganize their connectivity patterns, facilitating flexible and adaptive responses based on context, such as response to stimuli, circadian rhythmicity, or intrinsic states such as arousal and stress (Likhtik and Johansen 2019; McCormick et al. 2020; Zolin et al. 2021). Extensive research on neuromodulator actions has shown that their effects on circuit components can shape circuit output in a relatively consistent manner to meet these contextual demands (Nadim and Bucher 2014).

Here, we have proposed a somewhat different role for neuromodulation. Our findings suggest that convergent comodulation could have a significant impact on ensuring the consistent output of neural circuits, particularly when faced with substantial interindividual variability. Our findings do not contradict but rather complement the classical role for neuromodulators in providing flexibility of circuit output. For instance, even in the presence of multiple modulatory inputs, upregulation of a specific modulator could shift circuit output towards a modulator-specific pattern.

Modulatory actions clearly depend on the modulator concentrations and, in the case of modulatory neurotransmitters, on the cohort of co-transmitters as well as the specific targets of release (Nusbaum et al. 2001, 2017) which were not considered in the current study. At very low concentrations, threshold effects may in fact produce, rather than reduce, interindividual variability. The low concentration of 1 nM is close to threshold of *I*_MI_ activation for PROC and CCAP (Li et al. 2018). Since we kept the total modulator concentration constant, inconsistent changes in variability at low concentrations could be due to these threshold effects. At the other extreme, saturating concentrations may also have somewhat different effects on variability. Overall, the interindividual variability of the pyloric circuit output attributes were consistently lower at high peptide concentrations compared to mid concentrations (Fig. 9). However, comodulation at high concentrations did not consistently reduce variability compared to a single modulator. This is likely in part due to the saturation of neuropeptide receptors since receptor variability is also high across animals (Schneider et al. 2022). It therefore appears that the comodulatory actions that result in the reduction of interindividual variability occur at some goldilocks range of concentrations, perhaps consistent with tonic levels of these modulators in biological conditions.

### 7.4 Conclusions

The presence of interindividual variability of ion channel expression and synaptic properties poses the important question of how such variability may be constrained so that neural circuits could produce biologically meaningful output. The hypothesis that neuromodulators may be involved in compensating for interindividual variability well precedes the current study (Hamood and Marder 2014). We have modified this hypothesis to propose that it is the combined overlapping tonic presence of modulatory action that could provide a reasonable answer to constraining interindividual variability at the neural systems level. Considering that all neural systems are constantly targeted by actions of multiple modulators whose receptors have considerable overlapping subcellular actions (Doi and Ramirez 2008; Harris-Warrick 2011; Marder 2012; McCormick et al. 2020; Russo 2017), our findings justify the exploration of similar mechanisms in other neural circuits across animals.

## Acknowledgements

Current affiliation of ACS: University of Kassel, 34132 Kassel, Germany

Funding by NIH MH060605 to FN and DB, DFG SCHN 1594/1-1 to ACS

## 10 Extended Data legends

**Figure 2-1**

All data for the analysis shown in Figure 2D. The first column is the unique identifier for each experiment. A, B denotes the two preparations in the same dish from which we recorded simultaneously. Data columns show the average *F*_cycle_ in Hz over 30 s recordings (at least 10 bursts) for each experiment. Column headers indicate the experimental conditions in the order of application: intact, decentralized (neuromodulatory inputs removed), low (10^−9^ M) concentrations of PROC (P [low]), PROC + CCAP (PC [low]), PROC + CCAP + RPCH (PCR [low]), mid (3 x 10^−8^ M) concentrations of the same neuropeptide combinations (P [mid], PC [mid], PCR [mid]), wash. NaN indicates that the preparation was not rhythmic in that condition.

**Figure 3-1**

All data for the analysis shown in Figure 3. The Excel file contains three sheets, one for each neuron type (PD, LP, PY). The first column is the unique identifier for each experiment. A, B denotes the two preparations in the same dish from which we recorded simultaneously. Data columns show the average phase on and phase off over 30 s recordings (at least 10 bursts) for each experiment. Column headers indicate the experimental conditions in the order of application: intact, decentralized (neuromodulatory inputs removed), low (10^−9^ M) concentrations of PROC (P [low]), PROC + CCAP (PC [low]), PROC + CCAP + RPCH (PCR [low]), mid (3 x 10^−8^ M) concentrations of the same neuropeptide combinations (P [mid], PC [mid], PCR [mid]), wash. NaN indicates that the respective neuron was not active in that condition.

**Figure 4-1**

All data for the analysis shown in Figure 4. The Excel file contains three sheets, one for each neuron type (PD, LP, PY). The first column is the unique identifier for each experiment. A, B denotes the two preparations in the same dish from which we recorded simultaneously. Data columns show the average number of spikes (#spks) and spike frequency (Fspks) in Hz within a burst over 30 s recordings (at least 10 bursts) for each experiment. Column headers indicate the experimental conditions in the order of application: intact, decentralized (neuromodulatory inputs removed), low (10^−9^ M) concentrations of PROC (P [low]), PROC + CCAP (PC [low]), PROC + CCAP + RPCH (PCR [low]), mid (3 x 10^−8^ M) concentrations of the same neuropeptide combinations (P [mid], PC [mid], PCR [mid]), wash. NaN indicates that the respective neuron was not active or did not generate enough spikes to calculate spike frequency in that condition.

**Figure 5-1**

All data for the analysis shown in Figure 5. The Excel file contains four sheets, one for *F*_cycle_ and one for each neuron type (PD, LP, PY). In each sheet the first column is the unique identifier for each experiment. I, II denotes the two preparations in the same dish from which we recorded simultaneously. Data columns show the average *F*_cycle_ in Hz, phase on and off, number of spikes (#*spks*) and spike frequency (*F*_spks_) in Hz over 60 s recordings (at least 10 bursts) for each experiment. Column headers indicate the experimental conditions in the order of application: intact, decentralized (neuromodulatory inputs removed), mid (3 x 10^−8^ M) concentrations of PROC (P [mid]), PROC + CCAP (PC [mid]), PROC + CCAP + RPCH (PCR [mid]), wash. NaN indicates that the respective neuron or preparation was not active in that condition.

**Figure 6-1**

All data for the analysis shown in Figure 6C. The Excel file contains six sheets, the first three for the fit parameters (slope, curvature, I_0), four for hysteresis, and the last two are the raw f-I measurements for increasing (fI_incr.I) and decreasing (fI_decr.I) current injections. In each sheet the first column is the unique identifier for each experiment. Column headers indicate the experimental conditions in the order of application: intact, decentralized (neuromodulatory inputs removed), mid (3 x 10^−8^ M) concentrations of PROC (P [mid]), PROC + CCAP (PC [mid]), PROC + CCAP + RPCH (PCR [mid]), wash. The secondary column header in the last two sheets with the f-I measurements show the injected current in nA. NaN indicates that the respective neuron or preparation was not active in that condition. Empty cells indicate that that neuromodulator combination was not applied to those experiments.

**Figure 7-1**

All data for the analysis shown in Figure 7D. The Excel file contains four sheets, the first for the latency to the first spike in s, and the last three for the fit parameters (#spks_total, t_1/2, slope factor). In each sheet the first column is the unique identifier for each experiment. Column headers indicate the experimental conditions in the order of application: intact, decentralized (neuromodulatory inputs removed), mid (3 x 10^−8^ M) concentrations of PROC (P [mid]), PROC + CCAP (PC [mid]), PROC + CCAP + RPCH (PCR [mid]), wash. Latency is the latency (in s) over all five sweeps. NaN indicates that the respective neuron or preparation was not active in that condition. Empty cells indicate that that neuromodulator combination was not applied to those experiments.

**Figure 8-1**

All data for the analysis shown in Figure 8E. The Excel file contains four sheets, the first for the latency to the first spike in s, and the last three for the fit parameters (#spks_total, t_1/2, slope factor). In each sheet the first column is the unique identifier for each experiment. Column headers indicate the experimental conditions in the order of application: intact, decentralized (neuromodulatory inputs removed), mid (3 x 10^−8^ M) concentrations of PROC (P [mid]), PROC + CCAP (PC [mid]), PROC + CCAP + RPCH (PCR [mid]), wash. Latency is the latency (in s) over the last 10 of 20 sweeps. Fit parameters were obtained for the spike histograms of the last 10 of 20 sweeps. NaN indicates that the respective neuron or preparation was not active in that condition. Empty cells indicate that that neuromodulator combination was not applied to those experiments.

**Figure 9-1**

All data for the analysis shown in Figure 9. The Excel file contains four sheets, one for *F*_cycle_ and one for each neuron type (PD, LP, PY). In each sheet the first column is the unique identifier for each experiment. crab_1, crab_2 denotes the two preparations in the same dish from which we recorded simultaneously. Data columns show the average cycle frequency in Hz, phase on and off, number of spikes (#_spks_) and spike frequency (*F*_spks_) in Hz over 30 s recordings (at least 10 bursts) for each experiment. Column headers indicate the experimental conditions in the order of application: intact, decentralized (neuromodulatory inputs removed), mid (3 x 10^−8^ M) concentrations of PROC (P [mid]), PROC + CCAP (PC [mid]), PROC + CCAP + RPCH (PCR [mid]), wash. NaN indicates that the respective neuron or preparation was not active in that condition. Empty cells indicate that that neuromodulator combination was not applied to those experiments.

**Figure 10-1**

Model equations for the intrinsic and synaptic currents. The first column identifies the conductance, the second column the variables associated with that conductance, and the third column the equation for that parameter.

**Figure 10-2**

Parameters for the intrinsic currents. The first column denotes neuron, parameter, and current in the following form: neuron-parameter current. The second column is the value for that parameter.

**Figure 10-3**

Parameters for synaptic currents. The first column denotes pre- and postsynaptic neuron, synapse type, and parameter in the following form: presynaptic-postsynaptic synapsetype.parameter. The second column is the value for that parameter.

